# Structural and Mechanical Analysis of Treated and Untreated Aortic Coarctation in a Growing Porcine Model

**DOI:** 10.1101/2025.09.19.677411

**Authors:** Matthew A. Culver, Michael A. Stellon, Leah M. Gober, Sudhindra Chavadam, Dana Irrer, Luke Lamers, Alejandro Roldán-Alzate, Colleen M. Witzenburg

**Author notes:** Corresponding Author. Email address (C.M. Witzenburg).

## Abstract

Coarctation of the aorta (COA) is a congenital heart disease for which successful intervention can restore flow and reduce the blood pressure gradient, but does not ensure long-term health. Adults with successfully treated COA exhibit significantly higher incidence of hypertension. The objective of this study was to measure differences in the structure and mechanics of proximal and distal aortic tissue from a new, physiologically relevant growing porcine model of COA. This animal model also enabled the evaluation of a cutting-edge serially dilatable stent. Quantitative histologic analysis measured structural changes and the mechanical properties were investigated through uniaxial, shear lap, and peel tests of tissue from sham, control COA, and treated COA animals. Our original hypothesis that proximal aortic tissue from control and treated COA groups would be thicker and have less elastin was false. There were no significant differences in elastin content, collagen content, tissue area, lumen area, or lumen-to-tissue area between groups. Mechanically, distal tissue also exhibited no difference in either uniaxial or shear lap stiffness, failure stress, or failure strain between groups. Distal tissue from the COA control and treated COA groups however, exhibited, a lower circumferential failure peel tension, suggesting interlamellar strength was reduced. When compared with other previously published animal models of COA, a clear distinction was timing - our growing porcine model is the first for which COA was induced and treated at physiologically relevant time points. Our results indicated minimal adverse vascular remodeling in either the COA control or treated COA groups, however, it is unclear if this was due to a lack of severity, if elastinogenesis compensated for damage, or if another unknown mechanism prevented remodeling.

## INTRODUCTION

Coarctation of the aorta (COA) is a congenital heart disease (CHD) characterized by discrete narrowing of the aorta distal to the left subclavian artery. Each year, 1 in every 1,800 babies born in the United States exhibit COA, making it one of the most common CHDs with a prevalence ranging from 5-9% of all CHDs [1,2]. Severe forms of COA typically present early in life and successful early treatment has been linked to favorable long-term outcomes [3]. Left untreated, COA results in a significantly reduced life expectancy [4,5]. Indications for COA intervention include but are not limited to, a peak-to-peak systolic blood pressure gradient across the COA ≥ 20 mmHg, significant collateralization, and systemic ventricular dysfunction [6]. Surgery is the preferred form of treatment in small children and for more complex COA [7,8]. Currently, catheter-based interventions, including balloon angioplasty and stent implantation, are typically reserved for older patients and for post-surgical recurrent COA which occurs in approximately 10% of patients following surgical repair in infancy [2,9]. However, dilatable stents, which have recently received FDA approval for treatment of COA in infants, have been developed to adapt with the growing systemic vasculature [10]. Although endovascular treatment of COA has driven advances in this stent technology, trials designed to define superior therapeutic strategies remain conceptually remote, emphasizing the value of animal models for exploring key hemodynamic and physiological outcomes [11–13].

Surgical or catheter-based interventions can restore flow and reduce blood pressure gradients across the COA. However, *successful* intervention does not ensure long-term health. The prevalence of hypertension (HTN) in adults with treated COA is as high as 75% [14–19] and HTN is associated with increased risk of premature coronary artery disease, cerebrovascular incidents, heart failure, and aortic aneurysms [3,14–16,20–22]. Risk for post-stenotic dilatation and aneurysmal degeneration is also elevated following COA intervention [23–25]. The underlying cause of COA associated HTN and comorbidities is unclear. While various factors have been discussed, irreversible vascular remodeling has become a leading hypothesis. Histologic analysis of proximal aortic tissue from uncorrected COA patients is characterized by fibrosis, intimal thickening, and collagen and elastin degradation [26,27]. In addition, older case reports [28,29] indicate the histologic appearance of the elastic elements in the post-stenotic aortic wall from uncorrected COA patients were “defective” with focalized regions of “increased fragility.” Unfortunately, while pulse wave velocity and aortic distensibility (the two common clinical non-invasive methods to approximate aortic compliance) are significantly higher in hypertensive corrected COA patients [30,31], it is not possible to assess aortic remodeling non-invasively.

Animal models of COA have been developed in an attempt to characterize and quantify cardiovascular remodeling in response to disease and treatment. Increased wall thickness, reduced smooth muscle function, and elastin fragmentation were observed in the proximal aorta of both treated and untreated rabbit models of COA, suggesting adverse vascular remodeling persists following correction. In addition, thinning in the aorta distal to the COA region was reported. Although a successful model of COA, the age at which the rabbits underwent COA creation and treatment was not comparable to the typical timing in humans − a rabbit age of 10 weeks is approximately equivalent to a human age of 9 years [32].

Many previous studies have characterized the mechanical properties of healthy adult porcine aorta. For example, Witzenburg et al. and Shah et al. characterized the thoracic aorta from healthy pigs utilizing both uniaxial and biaxial testing [33,34]. Because of the complex layer structure of the aorta, multiple tests are required to capture rupture and dissection. In plane uniaxial extension tests characterize the stresses along the medial lamella which drive rupture [35–37]. However, peel and shear lap tests are necessary to capture the radial and shear stresses that drive the initiation and propagation of dissection [38–43]. Due to the increased risk of aortic aneurysms in patients with COA [44], it is crucial to implement each of these tests to fully capture aortic failure mechanics when evaluating tissue from COA animal models.

Irreversible adverse vascular remodeling has been implicated as a source of the long-term risk of recurrent hypertension in COA patients [14–16,18,45]. However, it is not possible to assess remodeling noninvasively and current animal models of COA induce stenosis at time points inconsistent with human disease. Our team created the first large animal model of COA in which stenosis was induced at an appropriate time-point and anatomic location and corrected at a time-point comparable to treatment of human disease [4]. Because we utilized a growing porcine model of COA, our team was also uniquely able to explore an early stent intervention with the ability for serial dilations throughout development to model treatment of COA [10]. The objective of this study was to test the hypotheses that 1) proximal aortic tissue from COA control and treated groups would exhibit thickening, reduced and or fragmented elastin, and higher stiffness in comparison to comparable tissue from the sham group and 2) distal tissue would exhibit a decrease in thickness, uniaxial and shear failure stresses, and peel tension. We evaluated remodeling of the aortic wall both proximal and distal to the COA site via quantitative histologic analysis and we characterized changes in the mechanical properties of the aortic wall distal to the COA via uniaxial, peel, and shear lap testing.

## 2. METHODS

### 2.1 ANIMAL MODEL

Fourteen domestic swine were obtained from the University of Wisconsin Swine Research and Teaching Center. In ten animals, discrete COA was surgically created in neonatal pigs at 2 weeks of age (4.5 ± 1.0 kg), which corresponds to ∼ 2 months of age in humans [4]. Six animals with COA underwent implantation of Renata MinimaTM (Renata Medical Company, Costa Mesa, CA) growth stent at 6 weeks of age. In these pigs stents were re-dilated to match somatic growth of the adjacent aorta at 12 and 20 weeks. Four additional animals served as sham controls. All groups had survival experimental catheterizations with angiographic imaging at 6 and 12 weeks and comprehensive terminal studies at 20 weeks (an age equivalent to adolescence in humans). Supplemental Figure 1 shows the experimental timeline. Within 3 hours of necropsy, the entirety of the aorta was collected from each animal for histologic analysis and mechanical testing. The Institutional Animal Care and Use Committee of the University of Wisconsin reviewed and approved this protocol.

#### Surgical COA creation

For a complete and detailed description of methods for surgical creation of COA see Stellon et al [4]. In brief, via a left thoracotomy, the proximal descending aorta was isolated and circumferentially mobilized. At this point, the sham group’s (n = 4) procedure was considered complete. For the COA animals (n = 10), the proximal descending aorta was cross-clamped and a 1 cm longitudinal incision was made in the left lateral side wall to induce intimal trauma. The arteriotomy was then plicated with a running 5-0 Prolene suture. A 5 mm long segment of 8 mm GORE-TEX tube graft was wrapped around the aorta and approximated with 5-0 Prolene suture to ensure external compression of the descending aortic diameter and localized stenosis over time.

### 2.2 HISTOLOGY

#### Collection

Histologic analysis was performed on regions of the aorta directly proximal and distal to the COA site for the COA control and treatment group as well as comparable regions for the sham group (Figure 1). At necropsy, these cross-sections were carefully extracted, fixed in 10% formalin for ∼24 hours, rinsed, placed in 70% ethanol, embedded in paraffin, and sectioned. Sections were stained with Masson’s trichrome (MTC) or Verhoeff–Van Gieson (VVG) to visualize collagen and elastin, respectively.

**Figure 1:**
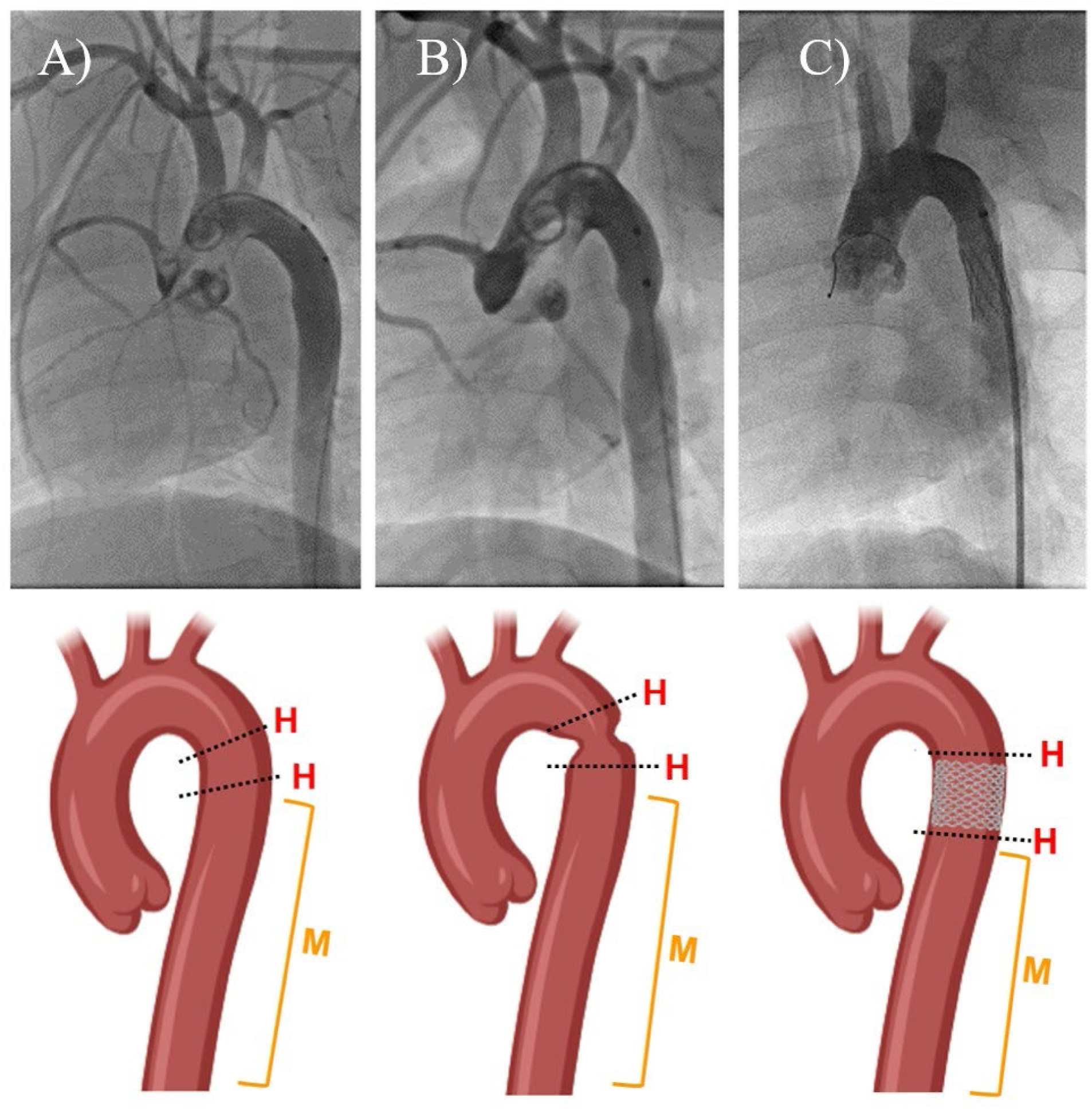
Angiograms and corresponding representative illustrations of the aorta of A) sham, B) COA, and C) treated groups. Dashed lines in illustrations marked ‘H’ were locations tissue cross-sections dissected for histologic analysis and the orange regions marked ‘M’ were locations tissue was dissected and tested mechanically.

#### Imaging

A Nikon Eclipse NiU microscope at a 10x objective was used to obtain images of each stained section. Due to the relatively large size of each section, the “manual large image grabber” feature was used to capture the entire aortic cross-section.

#### Segmentation

Two different researchers manually segmented VVG images to separate the media, intima, and lumen (Supplemental Figure 2) using Adobe Photoshop. First, they removed the image background and segmented the lumen using the color range tool. Next, they separated the elastin poor intima and adventitia from the elastin rich media by the color. The media, intima, and lumen were then mapped onto images of neighboring MTC stained slices. The researchers completed segmentation independently and were blinded to the group type, aorta location, and each other’s segmentation. Final measures of lumen and tissue area were normalized to animal weight.

#### Analysis

A custom MATLAB code developed by Bersi et al [46] was utilized to isolate the collagen and elastin components in the MTC and VVG stained segments, respectively, via colorimetric analysis (Figure 2). Utilizing predetermined hue, saturation, and lightness bounds corresponding to each stain, the number of pixels for each constituent was calculated and divided by the total number of colored pixels in the image to determine the area fraction of each constituent. Only the intima and media were included in the calculation of area fractions as the collagen-rich adventitia (Figure 2A) was often damaged during tissue collection.

**Figure 2:**
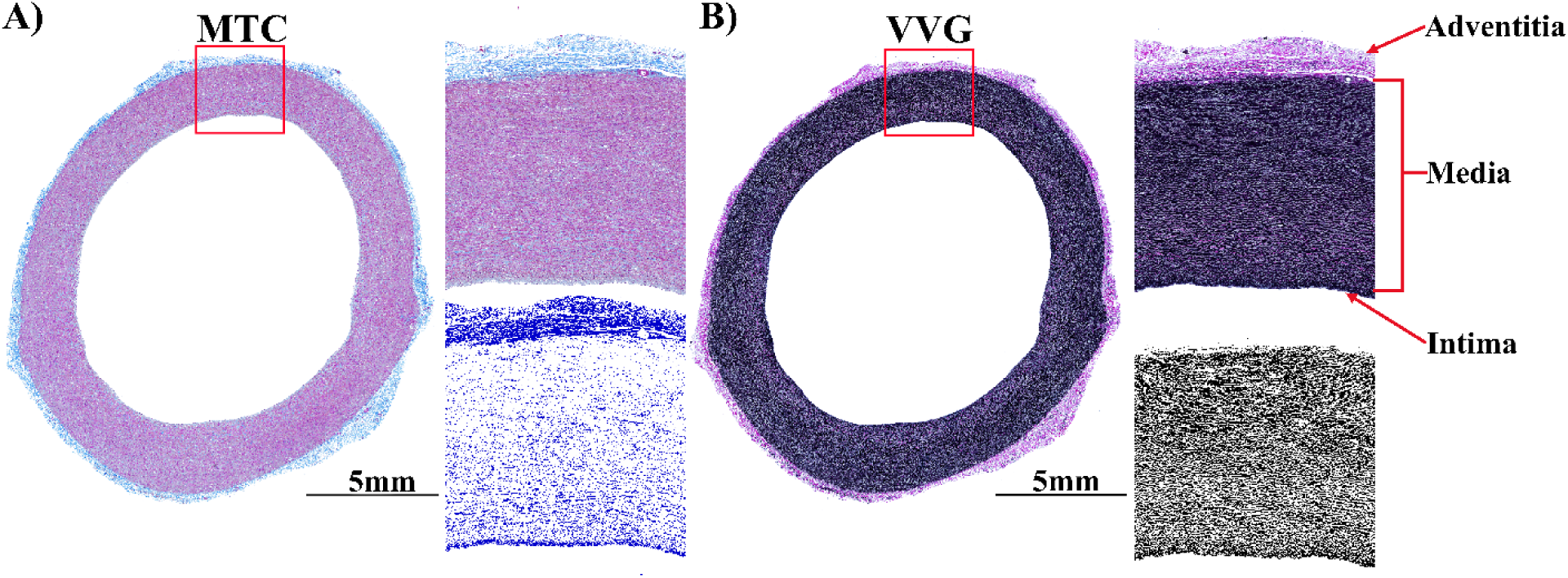
Representative images showing a A) Masson’s trichrome (MTC) and B) Verhoeff–Van Gieson (VVG) stained proximal sham aortic cross-section with automated visualization of collagen and elastin, respectively [46]. The collagen-rich adventitia was excluded from the final quantitative analysis.

### 2.3 MECHANICAL CHARACTERIZATION

#### Collection and Sample Preparation

Mechanical testing [33,34] was performed on descending aortic tissue directly distal to the COA site for the COA control and treatment groups as well as comparable regions for the sham group (Figure 1). At necropsy, the distal descending aorta was extracted and frozen at −80°C. Briefly, ∼24 hours after each descending aorta was removed from the freezer, a cut was made parallel to the long axis and the tissue was divided into 5×20 mm (uniaxial tests) or 5×10 mm (shear lap and peel tests) samples aligned either circumferentially or axially (Figure 3AB). Due to high incidence of slipping and clamp failure multiple samples were obtained in each direction from the descending aortic tissue directly distal to the COA site (within ∼ 20 mm) for each COA control and treatment animal or a comparable region for each sham animal. For uniaxial samples, 5mm biopsy punches were utilized to create a dog bone shape. For peel samples, a cut of approximately 3mm was made down the centerline, parallel to the plane of the aortic wall to initiate delamination. For shear lap samples, a cut of approximately 3mm was made on both ends centered within the medial layer and opposite sections were removed from each end to form an overlap region (Figure 3C). Previous studies indicate freezing at −80°C does not alter the mechanical behavior of the aorta [47–50].

**Figure 3:**
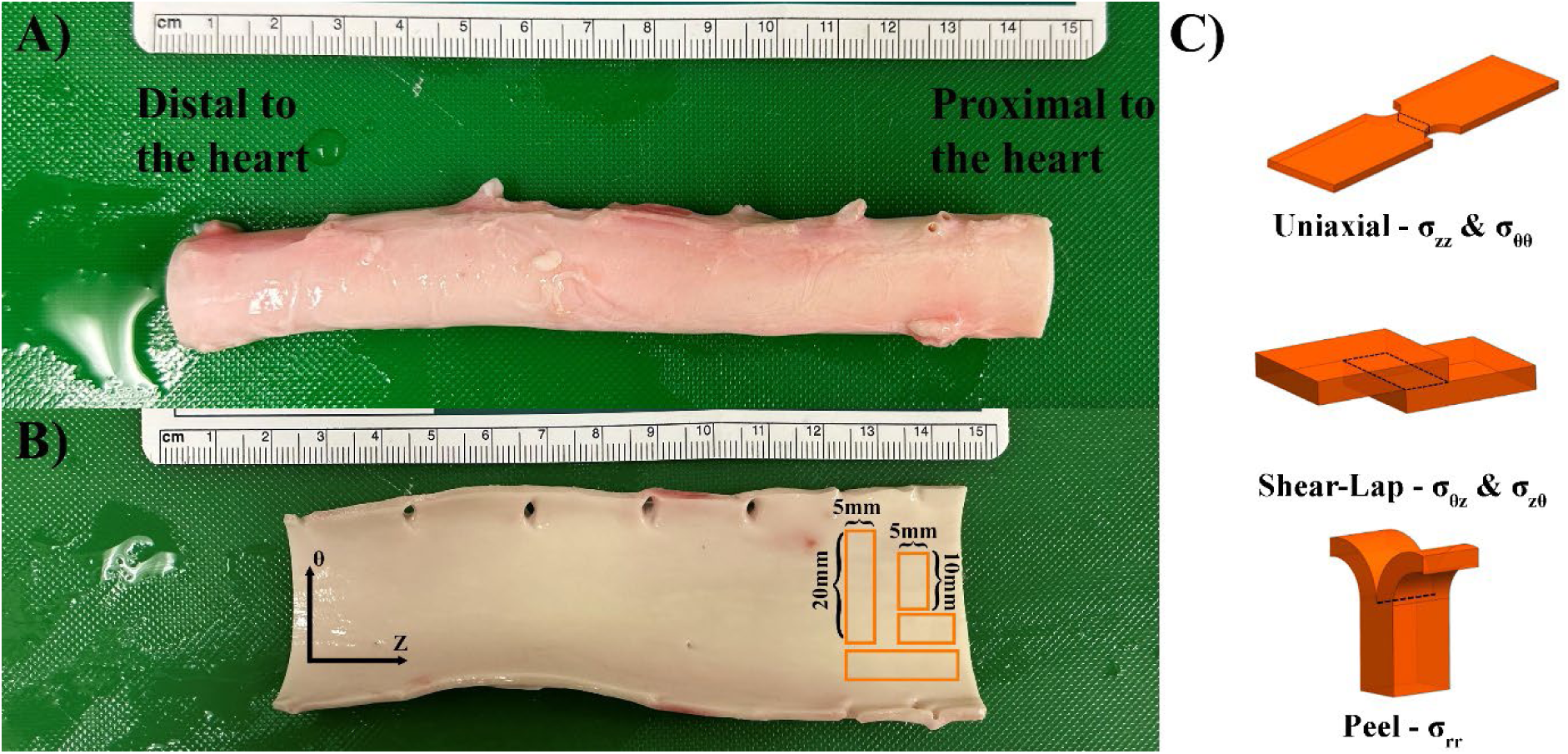
Specimen dissection for mechanical testing. The A) porcine descending aorta was B) cut open longitudinally along its medial edge and laid flat with the intimal surface up and the axial, Z, and circumferential, θ, axes along the horizontal and vertical directions, respectively. Rectangular sections were cut and shaped into C) uniaxial, peel, and shear lap samples. Pertinent areas of interest for computing normal and shear stress were shown on the idealized uniaxial and shear lap sample shapes, respectively (dashed rectangles). The pertinent width for computing peel tension was shown on the idealized peel sample shape (dashed line).

#### Testing

Uniaxial tests captured the tensile response along the axial and circumferential axes, peel tests captured the tensile response along the radial axis, and shear lap tests captured the response to shear along the medial lamella. Samples were clamped into a mechanical testing machine [51] (TestResources, E216SP Electro Dynamic Actuator) and pulled to failure to characterize their mechanical behavior at a rate of 3 mm/min [33,34]. Force was measured using load cells (TestResources, WF12S Miniature Fatigue Resistant Submersible IP65 100N) at a frequency of 100 Hz (TestResources, B8-16 TestBuilder and MTL32-2020). Images of each sample during testing were recorded at 6 Hz using a camera (Imperx, PoE-C2400, 2464 × 2056 pixels, 5 megapixels, 36 fps), lens (Computar, M3Z1228C-MP 0.66-Inch 1.3 Megapixel Varifocal lens 12-36mm F2.8 Manual Iris for uniaxial and peel tests and M3Z1228C-MP 0.66-inch 1.3 MP Varifocal Lens 12-36mm F2.8 Manual for shear lap tests), and image capture software (Imperx, IpxPlayer). The side of each shear lap sample was textured with charcoal powder to quantify full-field displacement. The side of each peel test was marked with at least two lines of India Ink to ensure delamination occurred along the sample centerline. Tests in which samples slipped or failed at the clamps were not included in the analysis.

#### Analysis

For the uniaxial samples we calculated the tensile first Piola-Kirchhoff stress, σ_T_, by dividing the grip force by the undeformed cross-sectional area in the gauge region (Figure 3C top). Stretch, λ, was calculated by dividing the recorded axial displacement by the initial gauge length for uniaxial tests. We initially utilized DIC derived Green-Lagrange tensile strain, but as the results did not differ substantially from the gauge length approach, we opted for the simpler method. Pre-failure stretch-stress data were fit with a piecewise constitutive model [52] such that

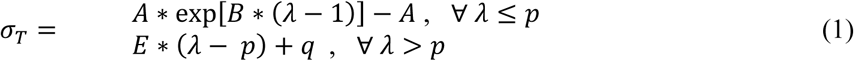

where *p* was the stretch at the transition point between the toe and linear regions of the stress-stretch curve. Similarly to Lee et al. [52], we used the MATLAB function fminsearch to simultaneously optimize *A* and *B* (the constants defining the stiffness of the toe region) as well as *p* by minimizing the mean summed squared difference in the model and measured stresses. The transition stress, *q*, and the stiffness of the linear region, *E*, were determined directly by enforcing continuity of the constitutive equation and its derivative at the transition point, respectively. Failure stress was defined as the stress at total or near total tissue separation with the corresponding stretch as the failure stretch. For shear lap tests, we calculated the shear first Piola-Kirchhoff stress, σ_*S*_, by dividing the grip force by the undeformed overlap area in the gage region (Figure 3C middle). Digital image correlation (DIC) was implemented to capture the full-field displacement and Green-Lagrange shear strain, γ [53]. Pre-failure strain-stress data were fit with a piecewise constitutive model such that

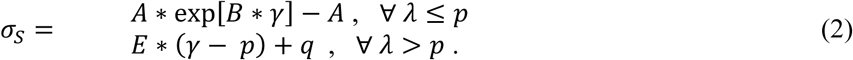

where *p* was the strain at the transition point between the toe and linear regions of the stress-stretch curve. Again, we used the MATLAB function fminsearch to simultaneously optimize *A*, *B*, and *p* by minimizing the mean summed squared difference stress and *q* and *E* were determined directly by enforcing continuity. Failure stress was defined as the stress at total or near total tissue separation with the corresponding strain as the failure strain. For peel tests, peel tension was calculated by dividing the grip force during peeling by the undeformed sample width (Figure 3C bottom). For each sample we reported the average peel tension since peeling occurred over an extended time frame. The initial positively sloped portion and final negatively sloped portion of the tension vs. time relationship were excluded since they indicate the portions of the test prior to the onset of peeling and during delamination completion, respectively (Figure 11A).

### 2.4 STATISTICAL ANALYSIS

Statistical analysis was performed in GraphPad Prism, version 10.1.2. Due to small group sizes, we employed the Kruskal-Wallis test with Dunn’s multiple comparisons, a conservative, non-parametric method, to determine if there were significant differences between groups. For histologic metrics a one sample Wilcoxon test determined if there were significant differences with location (i.e. proximal vs. distal to the coarctation site) for each group. For mechanical testing, multiple samples were obtained in each direction from the descending aortic tissue for each COA control and treatment animal or a comparable region for each sham animal. Therefore, for statistical analyses we first computed average metrics for samples from each pig. Then, we compared summary metrics between groups (sham, COA control, and treated). If mechanical comparisons via the Kruskal-Wallis test did not trend towards significance (defined as p < 0.1), measurements from the groups were pooled according to orientation and Wilcoxon matched-pairs signed rank test was performed to determine if there were significant differences for all animals between the axial and circumferential directions. Unless stated otherwise, p-values less than 0.05 were considered significant, p-values between 0.05 and 0.10 were considered trending towards significant, and summary values are reported as mean ± standard deviation.

## 3. RESULTS

### 3.1 ANIMAL MODEL

Figure 4 shows peak-to-peak systolic blood pressure gradient across the COA site for the COA control group (n = 4, 23.8 ± 11.6 mmHg) and across the same anatomic location for the sham group (n = 4, 0.3 ± 1.3) at 20 weeks. For the treated group (n = 6) the pressure gradient was measured before (8.7 ± 5.2 mmHg) and after (0.5 ± 2.3 mmHg) stent dilation. As expected, the pressure gradient was significantly higher in the COA control group than in the other groups. Twenty-week angiographic measurement of COA site diameter was also significantly different in the COA control group (6.4 ± 2.1 mm) than the treated COA group post-dilation (16.8 ± 1.5 mm) or the same location in the sham group (17.3 ± 1.0 mm) [4]. However, there were no statistically significant differences in ascending aortic systolic or diastolic pressure between groups (Supplemental Figure 3).

**Figure 4:**
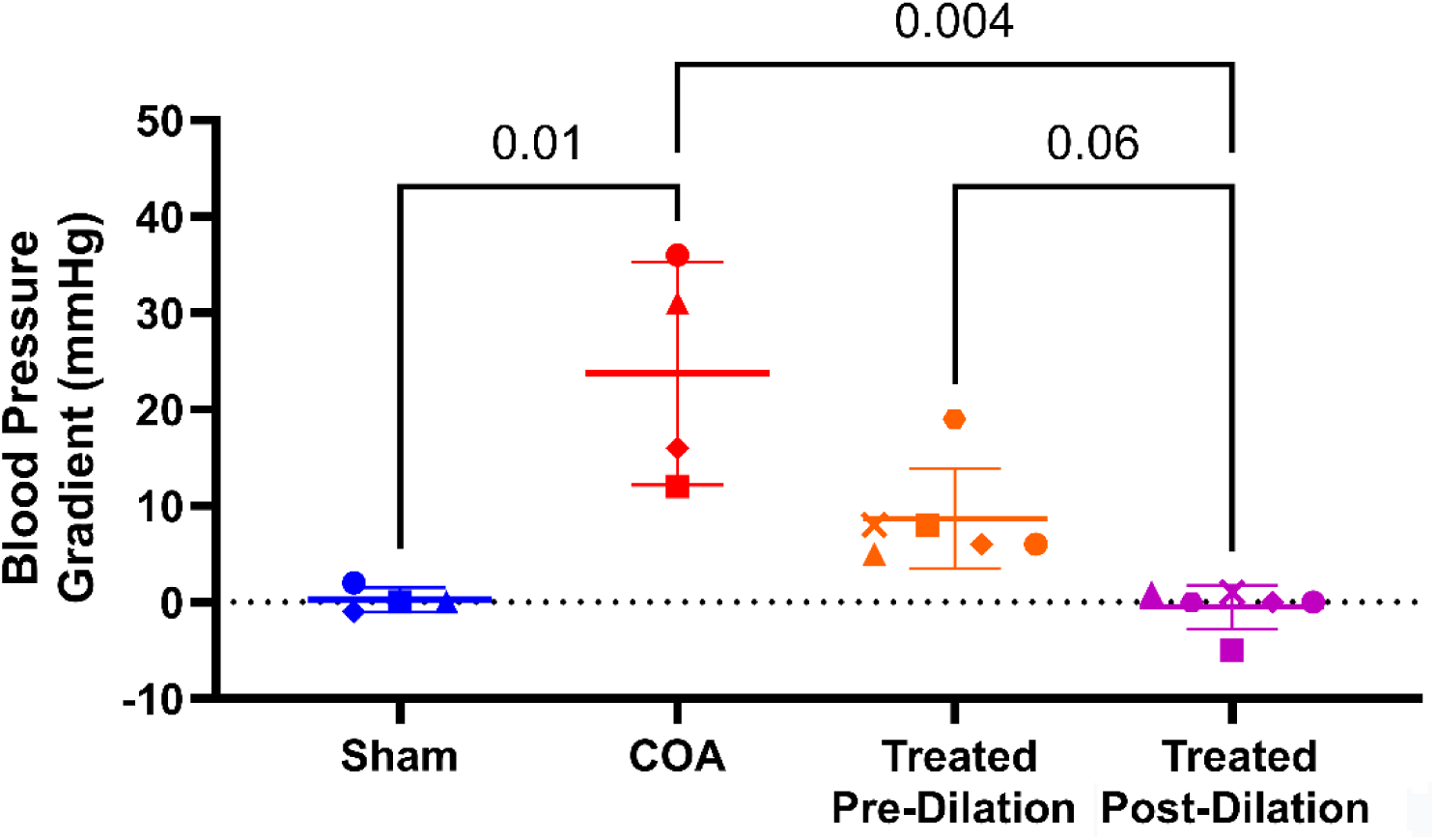
The peak-to-peak systolic blood pressure gradient across the COA site for the COA control group was significantly higher than the pressure gradient across the same anatomic location for the sham group at 20 weeks. It was also significantly higher than that of the treated group following stent dilation. Statistical significance was indicated for a Kruskal-Wallis test with Dunn’s multiple comparisons testing.

#### 3.2 HISTOLOGY

Figure 5 shows the elastin and collagen area fractions determined from colorimetric analysis of aortic cross-sections immediately proximal and distal to the COA site. Histologic analysis revealed trends toward greater elastin area fraction in the treated group compared to the sham and COA control groups both proximal and distal to the COA. However, this difference was only significant (or trending towards significance) in comparison to the COA group according to Dunn’s multiple comparison test following a Kruskal-Wallis test (Figure 5A). The locational change (i.e. proximal minus distal) in elastin area fraction was not significant across groups according to a Kruskal-Wallis test. According to a one sample Wilcoxon test this difference was significantly less than zero for the treated group. Therefore, treated group samples proximal to the COA site contained significantly less elastin than those taken distally. According to a Kruskal-Wallis test there were no significant differences between the three groups when comparing collagen area fraction for either the proximal or distal descending aorta (Figure 5C). Similarly, the locational change (i.e. proximal minus distal) in collagen area fraction was also not significant across groups according to a Kruskal-Wallis test. According to a one sample Wilcoxon test this difference was nearly significantly greater than zero for the treated group. Therefore, treated group samples proximal to the COA site contained more collagen than those taken distally.

**Figure 5:**
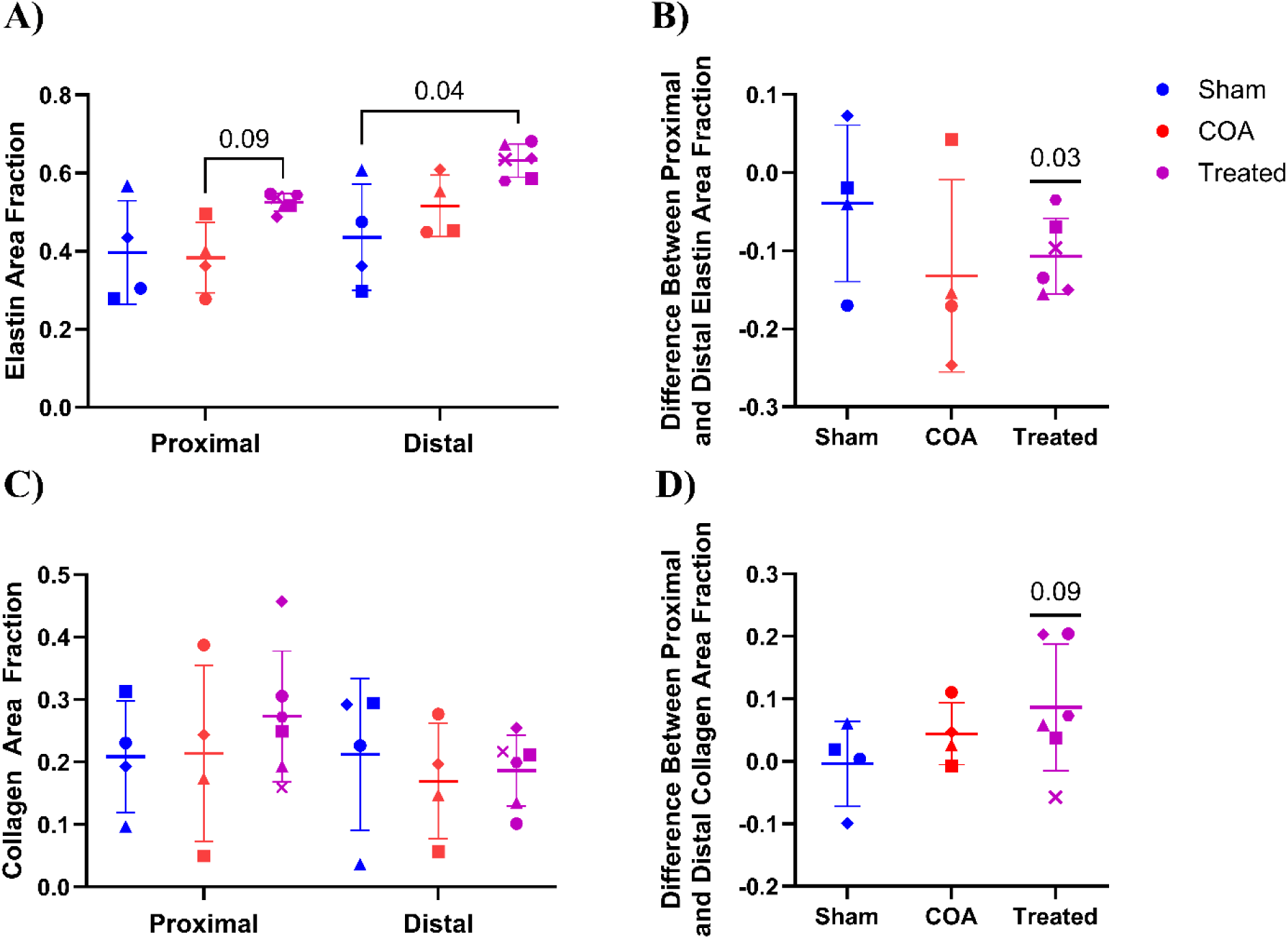
A) Elastin and C) collagen area fractions computed for aortic cross-sections located in the descending aorta immediately proximal and distal to the COA site from COA and treated animals as well as comparable locations for the sham animals. Treated group samples taken proximally to the COA site exhibited nearly significantly more elastin than proximal COA control samples and treated group samples taken distally to the COA site exhibited significantly more elastin than distal sham samples according to Dunn’s multiple comparison testing following a Kruskal-Wallis tests. There were no significant differences between groups in the location change (i.e. proximal minus distal) in B) elastin or D) collagen area fraction. However, a one sample Wilcoxon test indicated that in the treated animals there was a significantly lower elastin area fraction proximally and a nearly significantly higher collagen area fraction proximally.

We also compared the lumen and tissue (media + intima) areas as well as the ratio of lumen-to-tissue area to quantify vessel dilatation and thickening (Supplemental Figure 4 A-C). Lumen and tissue areas were normalized to animal weight, which did not differ significantly between groups but was variable (ranging from 61 to 94 kg). There were no significant differences between the three groups in lumen area, tissue area, or the ratio of lumen-to-tissue area. However, the proximal aorta of the treated group trended towards being thinner than that of the COA group according to the Dunn’s multiple comparisons test.

Qualitative assessment of histologic images revealed several structural findings. Untreated COA led to extensive collateralization to bypass the COA, consistent with presentation of chronic COA in humans [1,19]. Figure 6A shows an angiogram identifying a dilated internal mammary and collateral flow. Increased collateralization was not observed in the sham or treated COA groups at 20 weeks. Relative to their cross-sectional area, vessels branching from the internal mammary arteries (Figure 6B) were much thinner and had a larger ratio of lumen-to-tissue area than the descending aorta (Supplemental Figure 5) or carotid or femoral arteries (not shown) suggesting increased flow resulted in pathological remodeling. Marked intimal hyperplasia, characterized by pronounced irregular intimal thickening, was also observed in both the COA and treated groups (Supplemental Figure 5B and C) proximal and distal (not shown) to the COA site. Intimal hyperplasia was not observed in sham animals. Quantitatively, the proximal ratio of intima-to-tissue area was larger in both the COA control and treated groups than the sham group with the difference in the treated group nearing significance (Supplemental Figure 4D).

**Figure 6:**
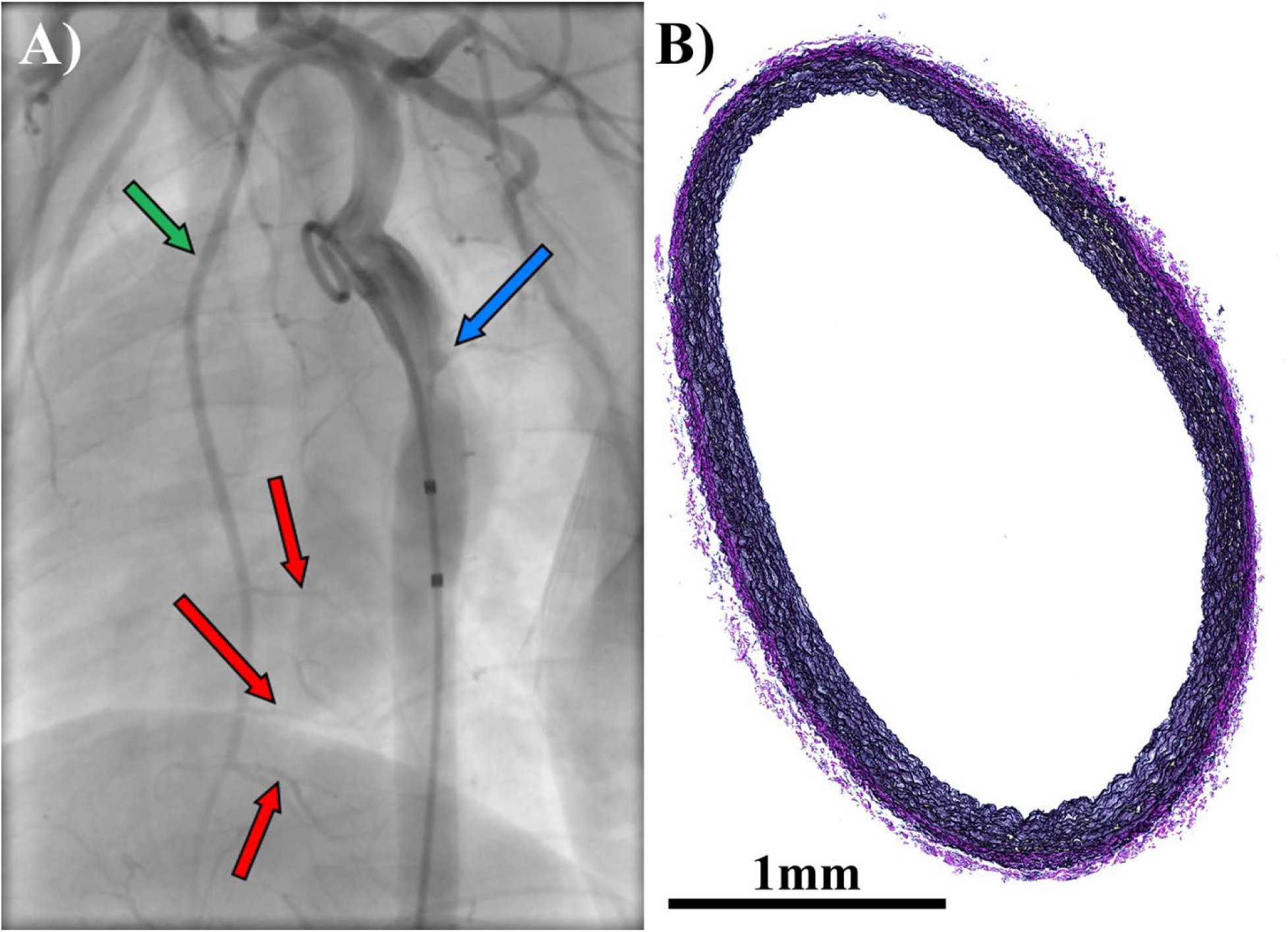
Representative A) angiogram and B) VVG stained section of collateral vessel from a COA animal. The blue arrow indicates COA site, the green arrow indicates a mammary artery, and the red arrows indicate collaterals sections taken for histology.

#### 3.3 MECHANICAL CHARACTERIZATION

##### Uniaxial Tests

Figure 7 shows the average first Piola-Kirchhoff tensile stress as a function of average grip stretch during uniaxial testing for samples from the three groups. Supplemental Figure 6A shows the constitutive model fit to representative sham samples in the axial and circumferential directions. In general, the model fit the data well and sum squared errors were less than 10% for 90% of the uniaxial samples. Neither *A* nor *B* (the toe region stiffness constants) differed between groups in either the axially or circumferentially aligned samples according to Kruskal-Wallis tests (Figure 8A). Similarly, *E* (the linear stiffness constant) did not differ between groups in either the axially or circumferentially aligned samples.

**Figure 7:**
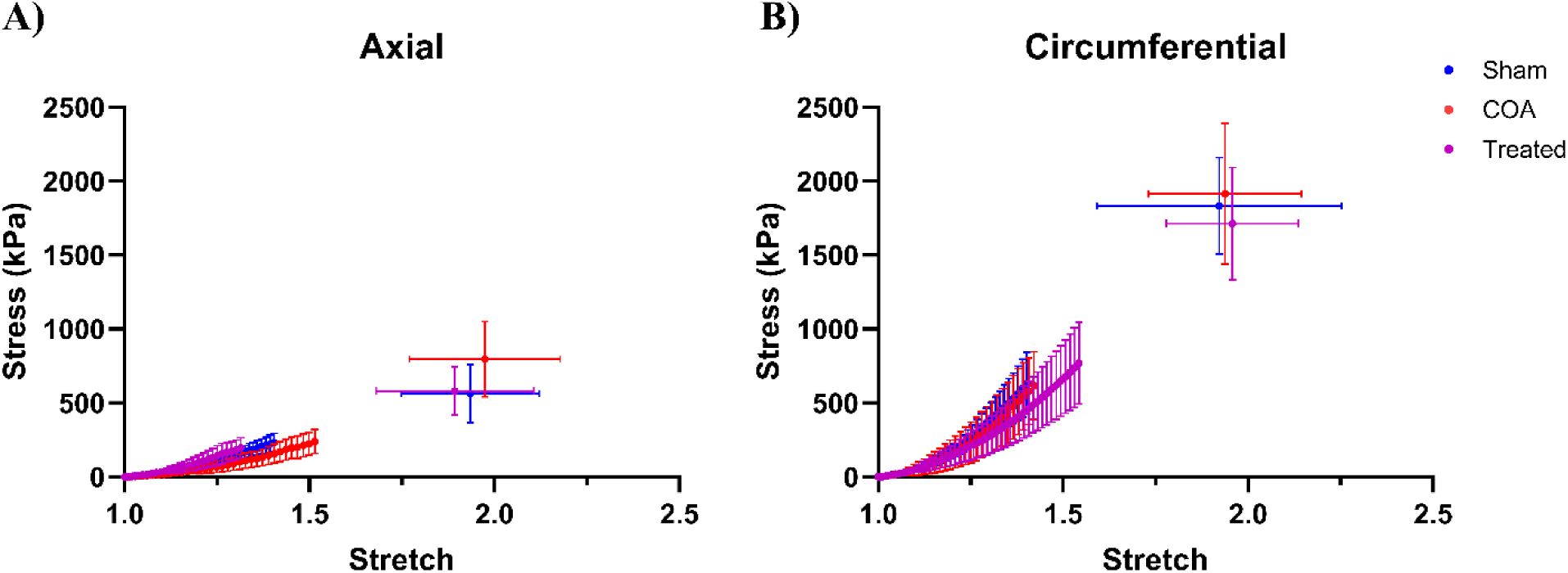
First Piola-Kirchhoff tensile (1^st^-PK) stress versus grip stretch during uniaxial extension to failure for samples oriented along the A) axial and B) circumferential directions (dots, mean +/- SD). Error bars are only shown for stretch up to the point at which the first sample failed. The final dot and error bars show the average stretch and stress at tissue failure +/- SD.

**Figure 8:**
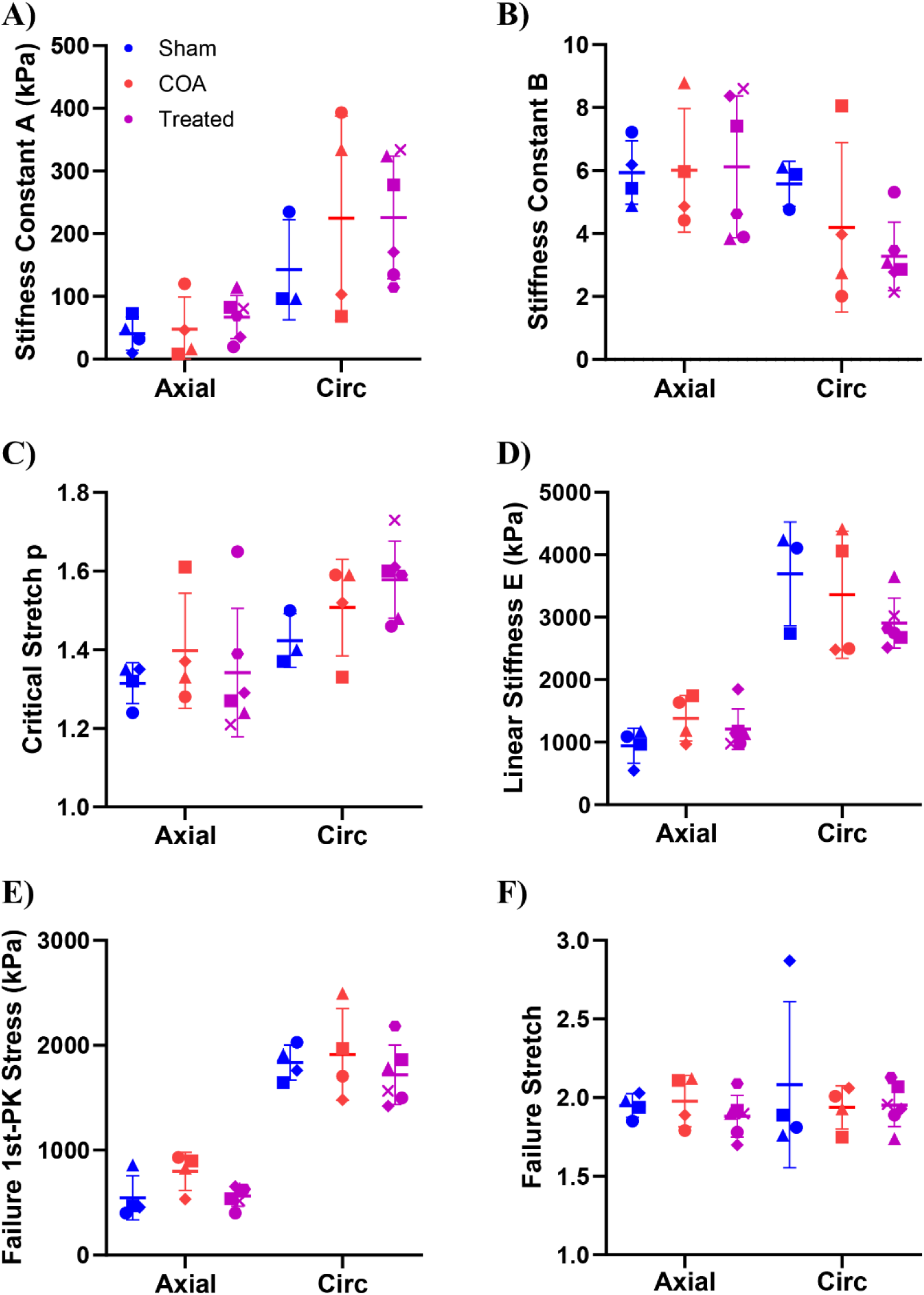
Fitted toe region stiffness constants A) *A* and B) *B* for uniaxial samples aligned axially (Axial) and circumferentially (Circ). C) The fitted transition stretch, *p*, between the toe and linear regions and D) the fitted linear stiffness constant, *E*. E) Failure first Piola-Kirchhoff tensile (1^st^-PK) stress and F) failure grip stretch. There were no statistically significant differences between groups in any metric according to Kruskal-Wallis tests (dots, mean +/- SD).

The average transition stretch, *p*, between the toe and linear regions was higher in the COA control and treated groups (Figure 8C; axial: COA 1.40 ± 0.15, treated 1.34 ± 0.16, circumferential: COA 1.51 ± 0.12, treated 1.58 ± 0.10) than the sham group (Figure 8C; axial: 1.32 ± 0.05, circumferential: 1.42 ± 0.07), however, this difference was not significant. Neither the failure tensile stress nor the failure stretch differed between groups in either the axially or circumferentially aligned samples. Supplemental Table 1 details the number of samples and animals included in the pre-failure and failure analysis for the uniaxial tests.

Next, we considered anisotropy in each metric by subtracting the axial value from the circumferential value. Anisotropy in *A*, *B*, *p*, *E*, failure tensile stress, and failure stretch did not differ significantly between groups according to Kruskal-Wallis tests (p-values ranged from 0.25 to 0.93). Therefore, data from the groups was pooled and a Wilcoxon matched-pairs signed rank test compared the axial and circumferential orientations (Supplemental Figure 7). *A*, *p*, *E*, and the failure stress were all significantly higher for circumferentially aligned samples than those aligned axially according to the Wilcoxon matched-pairs signed rank tests. In contrast *B* was higher in the axial direction, though this difference only approached significance.

##### Shear lap Tests

Figure 9 shows the average first Piola-Kirchhoff shear stress as a function of the average Green Lagrange shear strain during shear lap testing for samples from the three groups. Stresses were recorded for samples from all 14 pigs (4 sham, 4 COA, and 6 treated), however, some samples were excluded from shear strain calculations due to poor speckling (see Supplemental Table 1 for details). Supplemental Figure 6B shows the constitutive model fit to representative sham samples in the axial and circumferential directions. In general, the model fit the data well and sum squared errors were less than 10% for 90% of the shear lap samples. While *A* did not differ significantly between groups for the axially oriented samples (Figure 10A; sham 4.2 ± 1.9 kPa, COA 5.9 ± 6.8 kPa, treated 4.6 ± 3.0 kPa) it did for the circumferentially oriented samples (p-value = 0.033, sham 23.4 ± 15.7 kPa, COA 27.4 ± 0.8 kPa, treated 9.2 ± 4.6 kPa). The Dunn’s multiple comparisons test indicated a trend towards a lower stiffness constant in the treated group than the COA group, however, this is likely a result of our low animal numbers and the tight distribution of *A* for the COA group. Neither *B*, *p*, nor *E* differed between groups in either the axially or circumferentially aligned samples (Figure 10B, C, and D). Similarly, the failure shear stress and failure shear strain did not differ between groups in either the axially or circumferentially aligned samples (Figure 10E and F).

**Figure 9:**
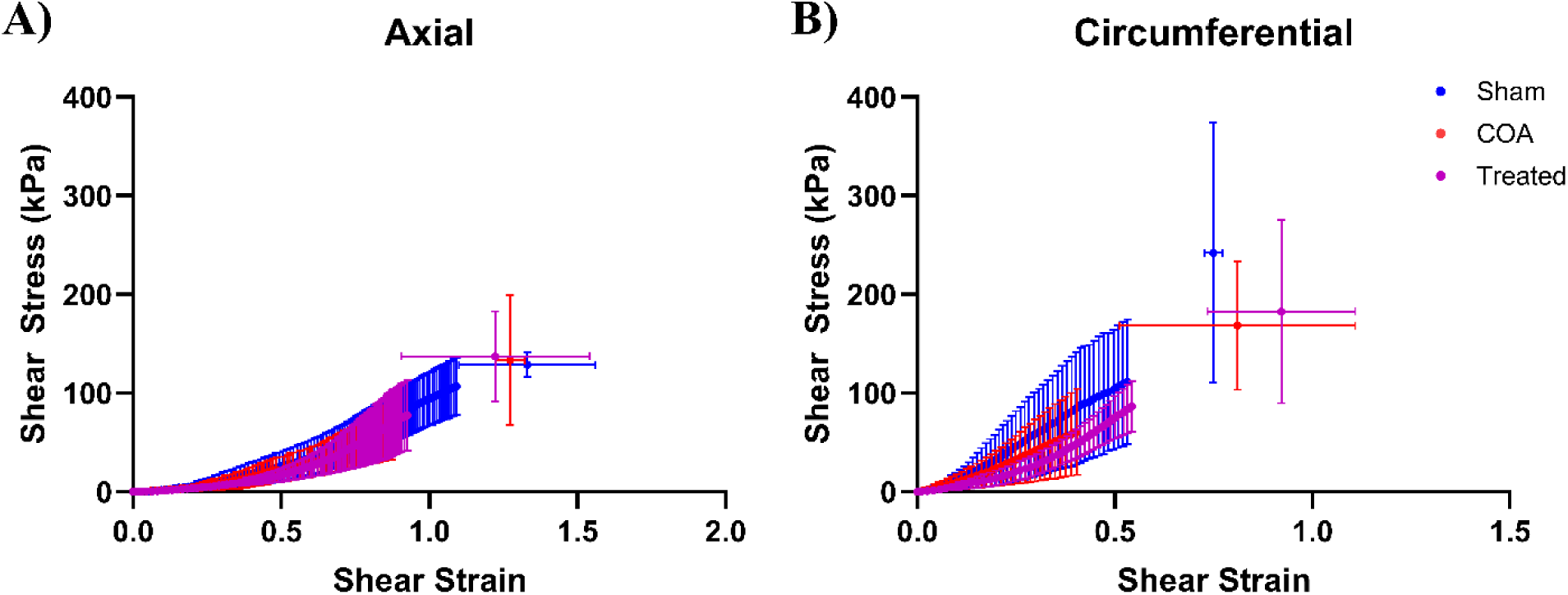
First Piola-Kirchhoff shear stress versus Green-Lagrange shear strain during shear lap extension to failure for samples oriented along the A) axial and B) circumferential directions (dots, mean +/- SD). Error bars are only shown for shear strain levels up to the point at which the first sample failed. The final dot and error bars shows the average shear strain and stress at tissue failure +/- SD.

**Figure 10:**
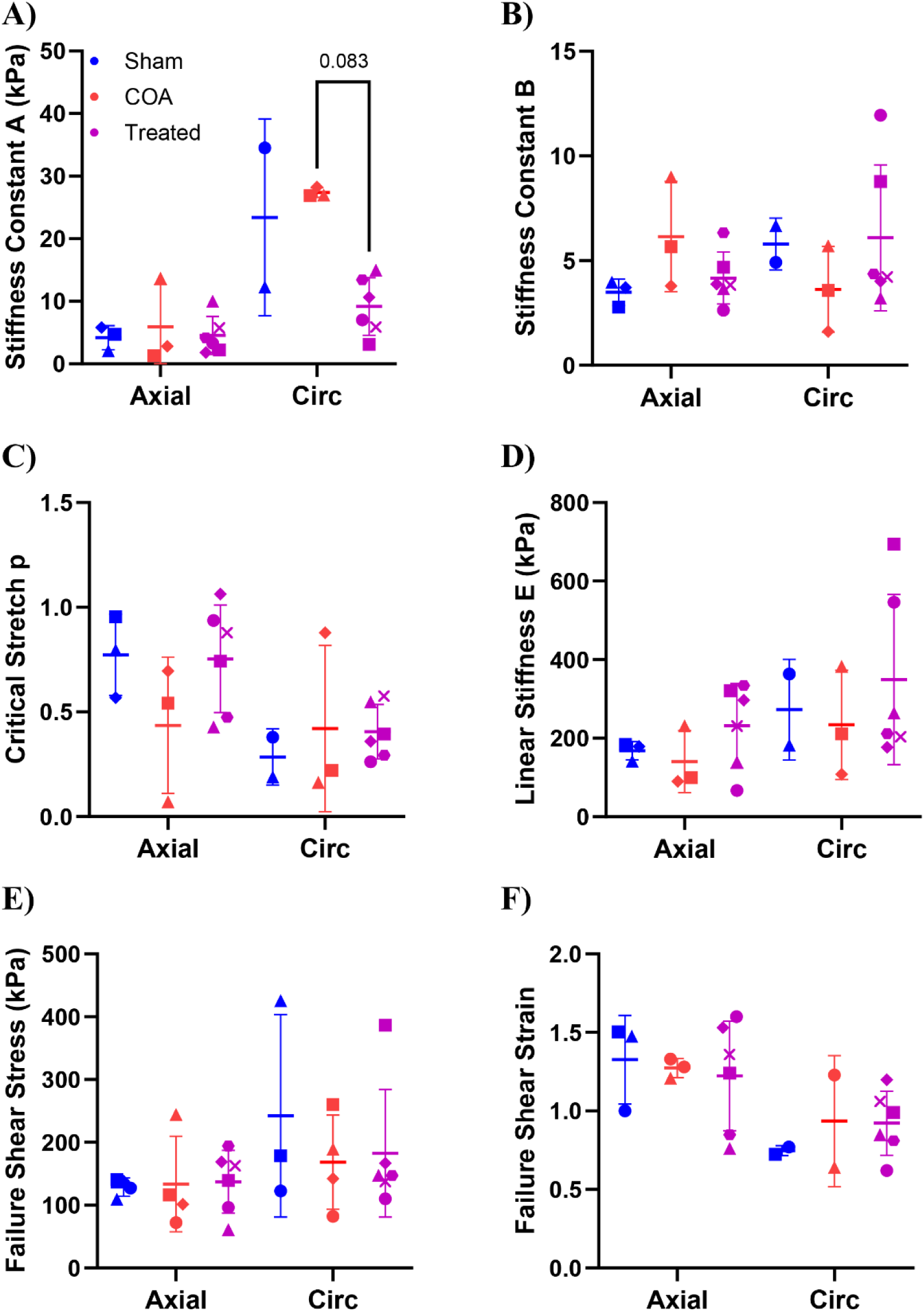
Fitted toe region stiffness constants A) *A* and B) *B* for shear lap samples aligned axially (Axial) and circumferentially (Circ). C) The fitted transition stretch, *p*, between the toe and linear regions and D) the fitted linear stiffness constant, *E*. E) Failure first Piola-Kirchhoff shear stress and F) failure Green-Lagrange shear strain. A Kruskal-Wallis test indicated *A* was significantly different between groups for circumferentially oriented samples with Dunn’s multiple comparisons testing indicating the difference between the COA and treated groups neared significance. There were no statistically significant differences between groups in any other metric (dots, mean +/- SD).

Next, we considered anisotropy in each metric by subtracting the axial value from the circumferential value. Anisotropy in *A* and *B* were significant between groups according to a Kruskal-Wallis test, therefore, data from the groups was not pooled. Instead, a Wilcoxon matched-pairs signed rank test comparing the axial and circumferential directions was performed on a group-wise basis. *A* was higher in the circumferentially aligned samples for the sham, COA, and treated groups (Δ*A* was 18 ± 17 kPa, 21 ± 7.1 kPa, and 4.6 ± 3.9 kPa, respectively). According to the Wilcoxon matched-pairs signed rank test this difference was significant for the treated group. In contrast, *B* was higher in the circumferentially aligned samples of the sham group (Δ*B* was 2.5 ± 0.6), but lower in the circumferentially aligned samples of the COA group (−2.5 ± 0.7), though neither of these differences were significant (Supplemental Figure 8AB). Anisotropy in *p*, *E*, failure shear stress, and failure Green-Lagrange shear strain did not differ significantly between groups according to a Kruskal-Wallis test (p-values ranged from 0.23 to 0.95). Therefore, data from the groups was pooled for each metric and a Wilcoxon matched-pairs signed rank test comparing the axial and circumferential orientations was performed (Supplemental Figure 8C-F). *p* and failure shear strain were significantly lower for circumferentially aligned samples than those aligned axially according to the Wilcoxon matched-pairs signed rank tests. In contrast, there were no significant differences in either *E* or failure shear stress between samples aligned circumferentially and axially.

##### Peel Tests

Figure 11 shows peel tension as a function of time for a representative sample (A) as well as the average peel tension (B) for samples from sham (4 pigs), COA (4 pigs), and treated groups (6 pigs). A Kruskal-Wallis test indicated no significant differences in average peel tension in the axial direction between groups. However, there was a nearly significant difference in average peel tension in the circumferential direction. The Dunn’s multiple comparisons test indicated a trend towards lower peel tension in both the COA and treated groups than the sham group for samples aligned circumferentially (Figure 11B; 28 ± 3 N/m, n = 9 and 29 ± 9 N/m, n = 15, respectively, vs. 43 ± 13 N/m, n = 8). Peel tension anisotropy (i.e. circumferential peel tension minus axial peel tension) was significant between groups according to a Kruskal-Wallis test. However, since there were nearly significant differences between groups in the circumferential direction data from the groups was not pooled. Instead, a Wilcoxon matched-pairs signed rank test comparing the axial and circumferential orientations was performed on a group-wise basis. The sham, COA, and treated groups exhibited lower average peel tensions in the circumferential direction than the axial direction (differences were −4.8 ± 29.1 N/m; −12.0 ± 2.8 N/m; −20.6 ± 11.2 N/m, respectively). According to the Wilcoxon matched-pairs signed rank test this difference was significant for the treated group.

**Figure 11:**
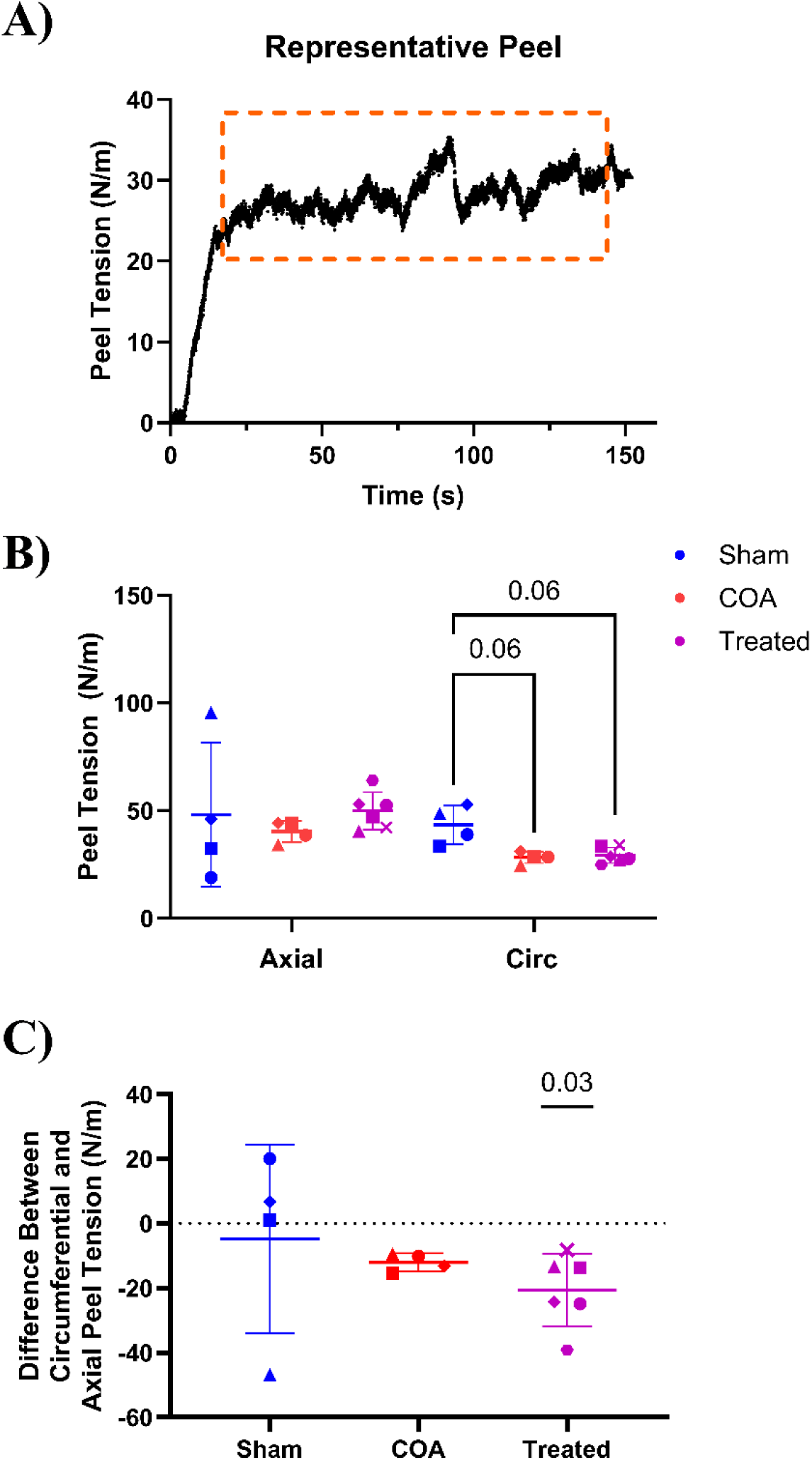
A) Representative peel tension as a function of time for a representative sample. B) Peel tension during failure for samples aligned axially (Axial) and circumferentially (Circ). Differences between groups neared significance for the circumferential direction according to a Kruskal-Wallis test with Dunn’s multiple comparison testing. C) Therefore, group-wise statistical significance between the Circ and Axial directions were indicated for Wilcoxon matched-pairs signed rank tests.

## 4. DISCUSSION

The objective of this study was to evaluate remodeling via quantitative histologic analysis and mechanical testing for a new, temporally relevant growing porcine model of treated and untreated COA. We hypothesized proximal aortic tissue from COA control and treated groups would be thicker, exhibit reduced and or fragmented elastin, and have a higher stiffness in comparison to the sham group. We also hypothesized distal aortic tissue from COA control and treated groups would exhibit a decrease in thickness, uniaxial and shear failure stresses, and peel tension. We evaluated remodeling of the aortic wall both proximal and distal to the COA site via quantitative histologic analysis and we characterized changes in the mechanical properties of the aorta wall distal to the COA via uniaxial, peel, and shear lap testing. Surprisingly, what we discovered was the COA control group exhibited no differences in aortic elastin or collagen content directly proximal or distal to the COA site when compared to the sham group. Intimal hyperplasia was observed in the control and treated COA groups but not in the sham group and extensive collateralization was present in untreated COA similar to what is seen in humans with untreated chronic moderate to severe COA. Additionally, the ratio of intimal-to-medial tissue area exhibited a trend towards significance in the treated COA group (Supplemental Figure 4D). Mechanically, there were no significant differences in either the uniaxial or shear failure stress or strain between the three groups. We did, however, observe a trend towards reduced peel tension in circumferentially aligned distal aortic samples from both the control and treated COA groups, potentially suggesting that radial strength was reduced.

### 4.1 HISTOLOGY

#### Proximal Aorta

In this model of COA created early in life in growing pigs, there were no significant differences in aortic elastin content (Figure 5A), collagen content (Figure 5B), lumen area (Supplemental Figure 4B), or lumen-to-tissue area (Supplemental Figure 4C) in tissue proximal to the COA site between the treated and untreated COA groups and the sham group. This is consistent with an adult mini-pig model of COA [54], but contrary to other COA studies. LaDisa and colleagues [32,55–57] reported a 2.8- and 2.0-fold increase in the tissue area of the aorta proximal to the COA site in untreated and treated rabbits, respectively, with no corresponding increase in elastin (leading to an overall reduction in the elastin area fraction). Hu et al. also reported lower elastin content and greater thickness of the proximal aorta in a mini-pig COA model in comparison to shams [58]. In contrast, Xu et al. reported conflicting results in their rabbit model of COA; with one study showing no difference in proximal aortic wall thickness or lumen area [59] and the other indicating increased proximal aortic wall thickness and lumen area [60]. A transient response in proximal aortic collagen content was observed in both mini-pig [58] and rabbit models of COA [59]. In adult mini-pigs quantitative histology indicated above-sham levels of collagen at 2 and 4 weeks following the creation of COA, but normal levels by weeks 6 and 8 [58]. Similarly, in adult rabbits, Western blot analysis indicated elevated levels of both type I and type III collagen and their procollagens at weeks 1, 2 and 4 following COA creation but sham levels by week 8 [59]. This transient increase is potentially consistent with our observations since animals were sacrificed at 20 weeks of age, a full 18 weeks after COA creation. In both rabbits [55,56] and mini-pigs [58] immunohistochemistry was also conducted to quantify smooth muscle content. There were no significant differences between sham and COA groups in smooth muscle α-actin content. However, Menon et al. reported higher levels of non-muscle myosin and lower levels of smooth muscle myosin in both the untreated and treated COA growing rabbits [55,56] and Hu et al. reported reduced caldesmon in untreated COA mini-pigs [58], suggesting a reduction in smooth muscle function. Additional experimental studies in growing pigs would be needed to assess smooth muscle content via immunohistochemistry.

#### Distal Aorta

In this growing porcine model of COA, there were also no significant differences in aortic elastin content (Figure 5A), collagen content (Figure 5B), tissue area (Supplemental Figure 4A), or lumen-to-tissue area (Supplemental Figure 4C) in tissue distal to the COA site between the control COA and sham groups. Notably, the elastin area fraction was significantly higher in the treated COA group than the control group. Stellon et al. [4] reported visible post-stenotic dilatation in both the control and treated COA animals, however, quantitative histology indicated that while lumen area distal to the COA site was greater in control COA animals than those in the treated or sham groups this difference was not significant (Supplemental Figure 4B) potentially suggesting in-vivo dilatation is a function of loading rather than an increase in circumferential tissue volume. Lin et al. [54,61] also conducted a quantitative histologic assessment of the post-stenotic region in their mini-pig model following COA creation. In their first study they reported significant dilatation in the post-stenotic region at 8 weeks [54], but in a later study they indicated that there was substantial individual variation, with 25% of COA animals exhibiting no post-stenotic dilatation at 12 weeks [61]. They also determined elastin content was significantly lower in post-stenotic tissue in comparison to a similar region for the sham group but there was no difference in either collagen content or smooth muscle α-actin content [54,61]. In contrast Menon et al. [55,56] reported no change in tissue area or elastin content distal to the COA, but they analyzed samples cut farther from the COA site (just proximal to the diaphragm) rather than in the post-stenotic region complicating comparison to both our results and those of Lin et al.

### 4.2 MECHANICAL CHARACTERIZATION

We observed differences between the control and treated COA groups and the sham group that neared significance in failure peel tension for circumferentially oriented distal aortic samples. Generally, peel testing estimates the interlamellar strength of a vessel, which is critical to aneurysm formation, dissection, and rupture [34,62]. To our knowledge there have been no peel tests on post-stenotic aortic tissue from animal models of COA. However, Korenczuk et al. completed peel testing on human ascending thoracic aortic aneurysm tissue resected during surgery [62]. They reported pre-failure behavior of aneurysmal tissue was statistically indistinguishable from previously published ascending thoracic porcine data, but that there were significant reductions in both circumferential and axial peel tension. Therefore, our findings may indicate a trend towards post-stenotic dysfunction. Yousef et al. recently reported that the ability of histologic microscopy to detect differences in the small, fine interlamellar structures of the aorta was inadequate and suggested higher resolution approaches [63]. Therefore, it is likely that our bulk measurements of elastin and collagen content from histologic imaging (Figure 5) were unable to detect interlamellar structural differences between groups. Serial block-face scanning electron microscopy has imaged the interlamellar structure of healthy rat abdominal aorta detailing the size and organization of the smooth muscle cells, the amount and orientation of interlamellar elastin fiber network, and the frequency of the sparse thick radial elastin struts which connect adjacent lamellae [64]. However, the time intensive nature and prohibitive cost of this approach limited this study to a single specimen.

Uniaxial and shear lap testing results did not differ between sham, control COA, and treated COA animals, but were consistent with previous mechanical tests on porcine ascending aorta. Uniaxial samples were significantly stiffer circumferentially than axially, exhibited much higher circumferential failure stresses, and similar failure strains [33,34]. The average pooled tensile failure stresses and stretches (625 ± 185 kPa and 1.93 ± 0.13 axially and 1809 ± 298 kPa and 1.99 ± 0.28 circumferentially) were remarkably similar to previously published values (753 ± 228 kPa and 1.91 ± 0.07 axially and 2510 ± 979 kPa and 1.99 ± 0.07 circumferentially) despite changes in the location along the aorta and animal source. Additionally, failure shear strains were lower circumferentially than axially, and failure shear stresses were higher [34]. The average pooled failure shear stresses (134 ± 49 kPa axially and 192 ± 104 kPa circumferentially) were also remarkably similar to previously published values (144 ± 16 kPa axially and 185 ± 28 kPa circumferentially) [34]. To our knowledge no other uniaxial or shear lap tests have been conducted on post-stenotic aortic tissue. Ghorbannia et al. conducted uniaxial testing of circumferentially oriented proximal aortic tissue from the growing rabbit model of aortic coarctation [65]. They reported increased stiffness with COA, however, this group also reported reduced elastin content and increased proximal aortic thickness in this same animal model [55,66].

### 4.3 SOURCES OF CONFLICT

Major differences in COA procedure and timing could have driven the differences in morphologic and immunohistochemical results observed between our growing porcine model and other previously published large animal models. Above all, the previously published models created COA in older animals. We created COA when the swine were only 2 weeks of age, which roughly corresponds to a 2-month-old human, to better replicate the typical clinical developmental stage at which COA is detected and treated [4]. LaDisa and colleagues created the only other model in which COA was induced prior to adulthood – the stenosis was generated at 10 weeks of age in the rabbits, corresponding roughly to 9 years of age in a human [32]. All other studies created the coarctation in adult animals [54,58–61]. Post-natal growth and development involves some level of elastinogenesis within the aorta, however, the extent and timing of this phase is unclear. Studies on the mouse aorta suggest the amount elastin doubles over the first 20 - 25 days of life (which is approximately equivalent to 4-5 months for a human infant [67]) but that the production rate then decays rapidly [68,69]. A study in rabbits indicates elastinogenesis ceases sometime between 6 and 10 weeks of age [70], suggesting no other animal study has induced or corrected COA during elastinogenesis. Therefore, one potential explanation for the lack of proximal aortic remodeling in the COA control group is that the stenosis was created early enough in development that the pigs could produce new elastin. This possibility was supported by the presence of elastin rich dilated internal mammary arteries off which collateral vessels arose in the COA control animals (Figure 6). Collateralization was not observed in other animal models [55,56], but is common in humans [71,72], further supporting the clinical relevance of our growing porcine model as more representative of human disease. It was also supported by the significantly higher level of elastin we observed distal to the COA site in the COA treated group.

Alternatively, the stenosis may have been less severe in our growing porcine model. COA duration in the swine (18 weeks) was longer or a similar time period as other animal models. Duration was 22 weeks in the growing rabbit model [32,55–57] and ranged from 8 to 12 weeks in the adult models [54,58–61]. However, the fixed nature of the stenosis in our pigs (∼ 5 mm diameter at the COA site) resulted in increasing COA severity as the animals grew. When the COA was induced in the 2-week-old swine, the diameter of the COA site was ∼ 2/3 of the same location in the sham group, whereas by the time the COA control swine were 20 weeks in age this ratio dropped to ∼1/3. Therefore, the pigs did not experience severe COA for the full 18 week follow-up period, and it is possible that remodeling may have been more severe had they lived longer or experienced a greater degree of stenosis earlier. The average peak-to-peak systolic blood pressure gradient in the growing pigs was > 20 mmHg at both 6 and 20 weeks for the control COA animals (Figure 4 and [4]), but the proximal aortic systolic and diastolic pressures were not statistically significantly higher in either the control or treated COA groups than the sham group at any time point (Supplemental Figure 3 and [4]). In the growing rabbit model [32], the post-surgical blood pressure gradient in the control COA group was 20 ± 5.3 mmHg (similar to the growing piglets), but the proximal mean blood pressure was 23 mmHg higher than the sham group. This was consistent with the minipig [58] model for which the proximal mean blood pressure was 20 mmHg higher in the COA group than the sham group.

Observations in the adult rabbit model more closely matched our growing porcine results, however, with a post-surgical blood pressure gradient between the ear and femoral arteries ranging from 15 to 31 mmHg and the mean arterial pressure at the ear artery just 4 mmHg larger than sham [59,60]. Similarly, in adult mini-pigs the average post-surgical proximal-to-distal blood pressure gradient was 33 vs. 6 mmHg in the COA group vs. the sham group, respectively, and the average mean proximal arterial pressure was slightly lower, 75 vs. 78 in COA group vs. the sham group, respectively [54]. Discrepancies in these models suggest differences in proximal mean aortic blood pressure may be an important metric to monitor in addition to the blood pressure gradient. There was also no significant difference in body weight in our study between the three groups at any age, which was consistent with other published studies [55,58–60]. Lastly, COA was surgically induced in the proximal descending aorta for our growing porcine model [4] and the published growing rabbit model [32,55–57]. Whereas, in adult models the stenosis was created at the mid-descending thoracic aorta [59,60], the suprarenal aorta [58], or infrarenal aorta [54,61], locations that are all inconsistent with typical COA presentation in humans.

### 4.4 LIMITATIONS AND FUTURE DIRECTIONS

To address the concern that stenosis may have been less severe initially in our growing porcine model than previous large animal models we surgically created COA in four additional animals. Terminal histologic staining and imaging were performed on regions of the aorta directly proximal and distal to the COA site for these animals as well as comparable regions for two additional sham animals. Unfortunately, catheterization measurements indicated that while the blood pressure gradient across the COA site was >20 mmHg it was not increased further (Supplemental Figure 9). While the addition of these animals did increase numerical power, addressing an additional limitation in this study, there was no change in overall results – elastin and collagen content did not differ between the sham and COA control group either proximally or distally (Supplemental Figure 10). It is possible additional animals would address the considerable variability observed in many of our measurements. Here, we relied on non-parametric statistical tests [73], taking a more conservative approach in our comparisons of groups. Further studies will employ the growing porcine COA model to characterize cardiovascular changes noninvasively via advanced MRI techniques. We will also conduct biochemical analysis, such as evaluating for the expression of endothelial C-type natriuretic peptide, to identify interventional therapeutic targets [74]. We are also seeking ways to quantify nascent elastin.

## 5. CONCLUSIONS

This growing porcine model of COA is advantageous over other animal models due largely to the physiologically relevant stage during which the stenosis is created. The comparable size and rapid growth of this animal model also enables evaluation of clinically relevant treatment options like the Renata Minima^TM^ growth stent. We evaluated proximal and distal aortic remodeling via quantitative histologic analysis and mechanical testing. Our original hypothesis that proximal aortic tissue from COA control and treated groups would exhibit reduced and or fragmented elastin and higher stiffness in comparison to the sham group was false in this animal model. However, it is unclear if the model severity or duration was insufficient to produce adverse proximal aortic remodeling, if elastinogenesis (which is supported by prominent collateralization and significantly higher distal elastin content in the stented group) compensated for this damage, or if another unknown mechanism prevented adverse remodeling. Overall, our results are consistent with clinical observations indicating repair at an earlier age is associated with better long-term health, including lower rates of chronic hypertension [19].

## ACKNOWLEDGMENTS

This study was funded by an award from the NIH-NCATS/UW Institute for Clinical and Translational Research (144-532905-AAK6656-4). The authors thank the University of Wisconsin Translational Research Initiatives in Pathology laboratory (TRIP) and the Experimental Animal Pathology Lab (EAPL), supported by the UW Department of Pathology and Laboratory Medicine, the University of Wisconsin Carbone Cancer Center (P30 CA014520), and the Office of The Director-NIH (S10 OD023526) for use of their facilities and services. The authors would also like to thank Michael Lasarev, MS for his help with statistical analysis.

## 6. AUTHOR STATEMENT

Matt A. Culver: Methodology, Validation, Formal analysis, Investigation, Writing - Original Draft, Visualization

Michael A. Stellon: Methodology, Formal analysis, and Investigation

Leah Gober: Validation, Formal analysis, Investigation, Data Curation, Writing – Review & Editing Sudhindra Chavadam: Validation, Formal analysis, Investigation

Dana Irrer: Conceptualization, Writing – Review & Editing, Supervision

Luke Lamers: Conceptualization, Methodology, Investigation, Resources, Data Curation, Writing – Review & Editing, Supervision, Project administration, Funding acquisition.

Alejandro Roldán-Alzate: Conceptualization, Methodology, Resources, Writing – Review & Editing, Supervision, Project administration, Funding acquisition.

Colleen M. Witzenburg: Conceptualization, Methodology, Formal analysis, Investigation, Resources, Writing - Original Draft, Writing – Review & Editing, Visualization, Supervision, Project administration, Funding acquisition.

## REFERENCES

[1] C.T. Mai, J.L. Isenburg, M.A. Canfield, R.E. Meyer, A. Correa, C.J. Alverson, P.J. Lupo, T. Riehle-Colarusso, S.J. Cho, D. Aggarwal, R.S. Kirby, National population-based estimates for major birth defects, 2010–2014, Birth Defects Res 111 (2019) 1420–1435. 10.1002/bdr2.1589.

[2] M.H.U. Usman, P. Rengifo-Moreno, S.F. Janzer, I. Inglessis-Azuaje, C. Witzke-Sanz, Coarctation of the Aorta: Management, Indications for Intervention, and Advances in Care, Curr Treat Options Cardiovasc Med 16 (2014). 10.1007/s11936-014-0341-2.

[3] E.J. Dijkema, T. Leiner, H.B. Grotenhuis, Diagnosis, imaging and clinical management of aortic coarctation, Heart 103 (2017) 1148–1155. 10.1136/heartjnl-2017-311173.

[4] M. Stellon, L. Gober, M.A. Culver, J. Hermsen, D. Irrer, C. Witzenburg, A. Roldán-Alzate, L. Lamers, Surgically induced aortic coarctation in a neonatal porcine model allows for longitudinal assessment of cardiovascular changes, Am J Physiol Heart Circ Physiol 326 (2024) H1117–H1123. 10.1152/ajpheart.00087.2024.

[5] M. Campbell, Natural history of coarctation of the aorta, Heart 32 (1970) 633–640. 10.1136/hrt.32.5.633.

[6] E.M. Isselbacher, O. Preventza, J. Hamilton Black, J.G. Augoustides, A.W. Beck, M.A. Bolen, A.C. Braverman, B.E. Bray, M.M. Brown-Zimmerman, E.P. Chen, T.J. Collins, A. DeAnda, C.L. Fanola, L.N. Girardi, C.W. Hicks, D.S. Hui, W. Schuyler Jones, V. Kalahasti, K.M. Kim, D.M. Milewicz, G.S. Oderich, L. Ogbechie, S.B. Promes, E. Gyang Ross, M.L. Schermerhorn, S. Singleton Times, E.E. Tseng, G.J. Wang, Y.J. Woo, D.P. Faxon, G.R. Upchurch, A.W. Aday, A. Azizzadeh, M. Boisen, B. Hawkins, C.M. Kramer, J.G.Y. Luc, T.E. MacGillivray, S.C. Malaisrie, K. Osteen, H.J. Patel, P.J. Patel, W.M. Popescu, E. Rodriguez, R. Sorber, P.S. Tsao, A. Santos Volgman, 2022 ACC/AHA Guideline for the Diagnosis and Management of Aortic Disease: A Report of the American Heart Association/American College of Cardiology Joint Committee on Clinical Practice Guidelines, Circulation 146 (2022). 10.1161/CIR.0000000000001106.

[7] A. Hager, C. Schreiber, S. Nützl, J. Hess, Mortality and Restenosis Rate of Surgical Coarctation Repair in Infancy: A Study of 191 Patients, Cardiology 112 (2009) 36–41. 10.1159/000137697.

[8] J.D.R. Thomson, A. Mulpur, R. Guerrero, Z. Nagy, J.L. Gibbs, K.G. Watterson, Outcome after extended arch repair for aortic coarctation., Heart 92 (2006) 90–4. 10.1136/hrt.2004.058685.

[9] M. Midulla, A. Dehaene, F. Godart, C. Lions, C. Decoene, W. Serge, M. Koussa, C. Rey, A. Prat, J.-P. Beregi, TEVAR in Patients With Late Complications of Aortic Coarctation Repair, Journal of Endovascular Therapy 15 (2008) 552–557. 10.1583/08-2436.1.

[10] E.M. Zahn, E. Abbott, N. Tailor, S. Sathanandam, D. Armer, Preliminary testing and evaluation of the renata minima stent, an infant stent capable of achieving adult dimensions, Catheterization and Cardiovascular Interventions 98 (2021) 117–127. 10.1002/ccd.29706.

11. B.H. Morray, K.F. Kennedy, D.B. McElhinney, Evolving Utilization of Covered Stents for Treatment of Aortic Coarctation: Report from the IMPACT Registry, Circ Cardiovasc Interv 16 (2023) E012697. 10.1161/CIRCINTERVENTIONS.122.012697.

[12] J.J.C. Gibb, W.C. Kim, F.G. Barlatay, A. Tometzki, A. Pateman, M. Caputo, D. Taliotis, Medium-Term Outcomes of Stent Therapy for Aortic Coarctation in Children Under 30 kg with New Generation Low-Profile Stents: A Follow-Up Study of a Single Centre Experience, Pediatr Cardiol 45 (2024) 544–551. 10.1007/s00246-023-03402-8.

[13] E. Bruckheimer, E. Birk, L. Benson, G. Butera, R. Martin, P.A. Roberts, M.B.E. Schneider, S. Schubert, H. Sievert, C.C.A. Pedra, Large Diameter Advanta V12 Covered Stent Trial for Coarctation of the Aorta: COARC Study, Circ Cardiovasc Interv 14 (2021) E010576. 10.1161/CIRCINTERVENTIONS.121.010576.

[14] D. Kenny, J.W. Polson, R.P. Martin, J.F.R. Paton, A.R. Wolf, Hypertension and coarctation of the aorta: An inevitable consequence of developmental pathophysiology, Hypertension Research 34 (2011) 543–547. 10.1038/hr.2011.22.

[15] S. Brili, D. Tousoulis, C. Antoniades, C. Aggeli, A. Roubelakis, S. Papathanasiu, C. Stefanadis, Evidence of vascular dysfunction in young patients with successfully repaired coarctation of aorta, Atherosclerosis 182 (2005) 97–103. 10.1016/j.atherosclerosis.2005.01.030.

[16] M. Heger, A. Willfort, T. Neunteufl, R. Rosenhek, H. Gabriel, G. Wollenek, M. Wimmer, G. Maurer, H. Baumgartner, Vascular dysfunction after coarctation repair is related to the age at surgery, Int J Cardiol 99 (2005) 295–299. 10.1016/j.ijcard.2004.02.001.

[17] S. Sendzikaite, R. Sudikiene, I. Lubaua, P. Silis, A. Rybak, G. Brzezinska-Rajszys, Ł. Obrycki, A. Jankauskiene, M. Litwin, Multi-centre cross-sectional study on vascular remodelling in children following successful coarctation correction, J Hum Hypertens 36 (2022) 819–825. 10.1038/s41371-021-00585-6.

[18] M.G.Y. Lee, R.A. Hemmes, J. Mynard, E. Lambert, G.A. Head, M.M.H. Cheung, I.E. Konstantinov, C.P. Brizard, G. Lambert, Y. d’Udekem, Elevated sympathetic activity, endothelial dysfunction, and late hypertension after repair of coarctation of the aorta, Int J Cardiol 243 (2017) 185–190. 10.1016/j.ijcard.2017.05.075.

[19] C. Canniffe, P. Ou, K. Walsh, D. Bonnet, D. Celermajer, Hypertension after repair of aortic coarctation - A systematic review, Int J Cardiol 167 (2013) 2456–2461. 10.1016/j.ijcard.2012.09.084.

[20] O.H. Toro-Salazar, J. Steinberger, W. Thomas, A.P. Rocchini, B. Carpenter, J.H. Moller, Long-Term Follow-Up of Patients After Coarctation of the Aorta Repair, 2002.

[21] P.M. Clarkson, M.R. Nicholson, B.G. Barratt-Boyes, J.M. Neutze, R.M. Whitlock, Results after repair of coarctation of the aorta beyond infancy: A 10 to 28 year follow-up with particular reference to late systemic hypertension, Am J Cardiol 51 (1983) 1481–1488. 10.1016/0002-9149(83)90661-6.

[22] M. Cohen, V. Fuster, P.M. Steele, D. Driscoll, D.C. McGoon, Coarctation of the aorta. Long-term follow-up and prediction of outcome after surgical correction., Circulation 80 (1989) 840–845. 10.1161/01.CIR.80.4.840.

[23] S. De Mey, P. Segers, I. Coomans, H. Verhaaren, P. Verdonck, Limitations of Doppler echocardiography for the post-operative evaluation of aortic coarctation, J Biomech 34 (2001) 951– 960. 10.1016/S0021-9290(01)00043-4.

[24] M. Campbell, J.H. Baylis, The Course and Prognosis of Coarctation of the Aorta, Heart 18 (1956) 475–495. 10.1136/hrt.18.4.475.

[25] X. Zhang, M. Luo, K. Fang, J. Li, Y. Peng, L. Zheng, C. Shu, Analysis of the formation mechanism and occurrence possibility of Post-Stenotic Dilatation of the aorta by CFD approach, Comput Methods Programs Biomed 194 (2020) 105522. 10.1016/j.cmpb.2020.105522.

26. J. Sehested, E. Mikkelsen, Different Reactivity and Structure of the Prestenotic and Poststenotic Aorta in Human Coarctation Implications for Baroreceptor Function, n.d. http://ahajournals.org.

[27] M.S. Dunnill, Histology of the aorta in coarctation., J Pathol Bacteriol 78 (1959) 203–7. http://www.ncbi.nlm.nih.gov/pubmed/13818687.

[28] F. Robicsek, P.W. Sanger, F.H. Taylor, R. Magistro, E. Foti, Pathogenesis and significance of post-stenotic dilatation in great vessels., Ann Surg 147 (1958) 835–844.

[29] H.K. de Vries, J.W. van den Berg, On the Origin of Poststenotic Dilatations, Cardiology 33 (1958) 195–211. 10.1159/000166341.

[30] E.J. Dijkema, M.G. Slieker, T. Leiner, H.B. Grotenhuis, Arterioventricular interaction after coarctation repair, Am Heart J 201 (2018) 49–53. 10.1016/j.ahj.2018.04.004.

31. A.L. Pauca, S.L. Wallenhaupt, N.D. Kon, Does Radial Artery Pressure Accurately Reflect Aortic Pressure?*, n.d.

[32] D.C. Wendell, I. Friehs, M.M. Samyn, L.M. Harmann, J.F. LaDisa, Treating a 20 mm Hg gradient alleviates myocardial hypertrophy in experimental aortic coarctation, Journal of Surgical Research 218 (2017) 194–201. 10.1016/j.jss.2017.05.053.

[33] S. Shah, C. Witzenburg, M.F. Hadi, H. Wagner, J. Goodrich, P. Alford, V.H. Barocas, Prefailure and Failure Mechanics of the Porcine Ascending Thoracic Aorta: Experiments and a Multiscale Model., J Biomech Eng 136 (2014) 4–10. http://www.ncbi.nlm.nih.gov/pubmed/24402447.

[34] C.M. Witzenburg, R.Y. Dhume, S.B. Shah, H.P. Wagner, J.A. Quindlen, P.W. Alford, V.H. Barocas, Failure of the Porcine Ascending Aorta: Multidirectional Experiments and a Unifying Model, J Biomech Eng 139 (2017) 031005–3. 10.1115/1.4035264.

[35] D. a Vorp, B.J. Schiro, M.P. Ehrlich, T.S. Juvonen, M.A. Ergin, B.P. Griffith, Effect of aneurysm on the tensile strength and biomechanical behavior of the ascending thoracic aorta., Ann Thorac Surg 75 (2003) 1210–4.

[36] D.C. Iliopoulos, E.P. Kritharis, A.T. Giagini, S.A. Papadodima, D.P. Sokolis, Ascending thoracic aortic aneurysms are associated with compositional remodeling and vessel stiffening but not weakening in age-matched subjects, Journal of Thoracic and Cardiovascular Surgery 137 (2009) 101–109. 10.1016/j.jtcvs.2008.07.023.

[37] J.E. Pichamuthu, J.A. Phillippi, D.A. Cleary, D.W. Chew, J. Hempel, D.A. Vorp, T.G. Gleason, Differential tensile strength and collagen composition in ascending aortic aneurysms by aortic valve phenotype, Annals of Thoracic Surgery 96 (2013) 2147–2154. 10.1016/j.athoracsur.2013.07.001.

[38] C. van Baardwijk, M.R. Roach, Factors in the propagation of aortic dissections in canine thoracic aortas., J Biomech 20 (1987) 67–73.

[39] G. Sommer, T.C. Gasser, P. Regitnig, M. Auer, G. a Holzapfel, Dissection properties of the human aortic media: an experimental study., J Biomech Eng 130 (2008) 021007. 10.1115/1.2898733.

[40] J. Tong, G. Sommer, P. Regitnig, G. a Holzapfel, Dissection properties and mechanical strength of tissue components in human carotid bifurcations., Ann Biomed Eng 39 (2011) 1703–19. 10.1007/s10439-011-0264-y.

[41] A. Tsamis, S. Pal, J.A. Phillippi, T.G. Gleason, S. Maiti, D.A. Vorp, Effect of aneurysm on biomechanical properties of “radially-oriented” collagen fibers in human ascending thoracic aortic media, J Biomech 47 (2014) 3820–3824. 10.1016/j.jbiomech.2014.10.024.

[42] M. Kozuń, Delamination properties of the human thoracic arterial wall with early stage of atherosclerosis lesions, Journal of Theoretical and Applied Mechanics (Poland) 54 (2016) 229–238. 10.15632/jtam-pl.54.1.229.

[43] D.E. Gregory, J.H. Veldhuis, C. Horst, G. Wayne Brodland, J.P. Callaghan, Novel lap test determines the mechanics of delamination between annular lamellae of the intervertebral disc., J Biomech 44 (2011) 97–102. 10.1016/j.jbiomech.2010.08.031.

[44] S. Raza, S. Aggarwal, P. Jenkins, A. Kharabish, S. Anwer, D. Cullington, J. Jones, J. Dua, V. Papaioannou, R. Ashrafi, S. Moharem-Elgamal, Coarctation of the Aorta: Diagnosis and Management, Diagnostics 13 (2023). 10.3390/diagnostics13132189.

[45] R.M. Radke, G.-P. Diller, M. Duck, S. Orwat, D. Hartmann, T. Thum, H. Baumgartner, Endothelial function in contemporary patients with repaired coarctation of aorta, Heart 100 (2014) 1696–1701. 10.1136/heartjnl-2014-305739.

[46] M.R. Bersi, C. Bellini, J. Wu, K.R.C. Montaniel, D.G. Harrison, J.D. Humphrey, Excessive adventitial remodeling leads to early aortic maladaptation in angiotensin-induced hypertension, Hypertension 67 (2016) 890–896. 10.1161/HYPERTENSIONAHA.115.06262.

[47] S.A. O’Leary, B.J. Doyle, T.M. McGloughlin, The impact of long term freezing on the mechanical properties of porcine aortic tissue, J Mech Behav Biomed Mater 37 (2014) 165–173. 10.1016/j.jmbbm.2014.04.015.

[48] B.D. Stemper, N. Yoganandan, M.R. Stineman, T.A. Gennarelli, J.L. Baisden, F.A. Pintar, Mechanics of Fresh, Refrigerated, and Frozen Arterial Tissue, Journal of Surgical Research 139 (2007) 236–242. 10.1016/j.jss.2006.09.001.

[49] J.O. Virues Delgadillo, S. Delorme, R. El-Ayoubi, R. DiRaddo, S.G. Hatzikiriakos, Effect of freezing on the passive mechanical properties of arterial samples, J Biomed Sci Eng 03 (2010) 645– 652. 10.4236/jbise.2010.37088.

[50] A. Hemmasizadeh, K. Darvish, M. Autieri, Characterization of changes to the mechanical properties of arteries due to cold storage using nanoindentation tests, Ann Biomed Eng 40 (2012) 1434–1442. 10.1007/s10439-011-0506-z.

[51] D. Pearce, M. Nemcek, C. Witzenburg, Combining Unique Planar Biaxial Testing with Full-Field Thickness and Displacement Measurement for Spatial Characterization of Soft Tissues, Curr Protoc 2 (2022). 10.1002/cpz1.493.

[52] A.H. Lee, S.E. Szczesny, M.H. Santare, D.M. Elliott, Investigating mechanisms of tendon damage by measuring multi-scale recovery following tensile loading, Acta Biomater 57 (2017) 363–372. 10.1016/j.actbio.2017.04.011.

[53] R. Rahupathy, V. Barocas, Robust Image Correlation Based Strain Calculator for Tissue Systems (20130022, Dr. Victor Barocas), https://license.umn.edu/product/robust-image-correlation-based-strain-calculator-for-tissue-systems (2013).

[54] P.Y. Lin, Y.T. Wu, G.C. Lin, Y.H. Shih, A. Sampilvanjil, L.R. Chen, Y.J. Yang, H.L. Wu, M.J. Jiang, Coarctation-induced degenerative abdominal aortic aneurysm in a porcine model, J Vasc Surg 57 (2013). 10.1016/j.jvs.2012.08.104.

[55] A. Menon, D.C. Wendell, H. Wang, T.J. Eddinger, J.M. Toth, R.J. Dholakia, P.M. Larsen, E.S. Jensen, J.F. LaDisa, A coupled experimental and computational approach to quantify deleterious hemodynamics, vascular alterations, and mechanisms of long-term morbidity in response to aortic coarctation, J Pharmacol Toxicol Methods 65 (2012) 18–28. 10.1016/j.vascn.2011.10.003.

[56] A. Menon, T.J. Eddinger, H. Wang, D.C. Wendell, J.M. Toth, J.F. LaDisa, Altered hemodynamics, endothelial function, and protein expression occur with aortic coarctation and persist after repair, Am J Physiol Heart Circ Physiol 303 (2012) 1304–1318. 10.1152/ajpheart.00420.2012.-Coarctation.

[57] J. Azarnoosh, A. Ghorbannia, E.S.H. Ibrahim, H. Jurkiewicz, L. Kalvin, J.F. LaDisa, Temporal evolution of mechanical stimuli from vascular remodeling in response to the severity and duration of aortic coarctation in a preclinical model, Sci Rep 13 (2023). 10.1038/s41598-023-34400-8.

[58] J.J. Hu, A. Ambrus, T.W. Fossum, M.W. Miller, J.D. Humphrey, E. Wilson, Time courses of growth and remodeling of porcine aortic media during hypertension: A quantitative immunohistochemical examination, Journal of Histochemistry and Cytochemistry 56 (2008) 359–370. 10.1369/jhc.7A7324.2007.

[59] C. Xu, C.K. Zarins, H.S. Bassiouny, W.H. Briggs, C. Reardon, S. Glagov, Differential Transmural Distribution of Gene Expression for Collagen Types I and III Proximal to Aortic Coarctation in the Rabbit, J Vasc Res 37 (2000) 170–182. 10.1159/000025728.

[60] C. Xu, C.K. Zarins, S. Glagov, Gene expression of tropoelastin is enhanced in the aorta proximal to the coarctation in rabbits, Exp Mol Pathol 72 (2002) 115–123. 10.1006/exmp.2002.2423.

[61] B.W. Lin, P.Y. Lin, Y.H. Shih, C.L. Wu, Y.T. Wu, M.J. Jiang, High-Frequency Fluctuation Intensity as a Predictor of Post-Stenotic Dilatation in Swine, IEEE Trans Biomed Eng 70 (2023) 991–999. 10.1109/TBME.2022.3207060.

[62] C.E. Korenczuk, K.K. Liao, V.H. Barocas, Ex Vivo Mechanical Tests and Multiscale Computational Modeling Highlight the Importance of Intramural Shear Stress in Ascending Thoracic Aortic Aneurysms, 141 (2019) 1–11. 10.1115/1.4045270.

[63] S. Yousef, N. Matsumoto, I. Dabe, M. Mori, A.B. Landry, S.R. Lee, Y. Kawamura, C. Yang, G. Li, R. Assi, P. Vallabhajosyula, A. Geirsson, G. Moeckel, J.D. Humphrey, G. Tellides, Quantitative not qualitative histology differentiates aneurysmal from nondilated ascending aortas and reveals a net gain of medial components, Sci Rep 11 (2021) 1–13. 10.1038/s41598-021-92659-1.

[64] M.K. O’Connell, S. Murthy, S. Phan, C. Xu, J.A. Buchanan, R. Spilker, R.L. Dalman, C.K. Zarins, W. Denk, C.A. Taylor, The three-dimensional micro- and nanostructure of the aortic medial lamellar unit measured using 3D confocal and electron microscopy imaging, Matrix Biology 27 (2008) 171–181. 10.1016/j.matbio.2007.10.008.

[65] A. Ghorbannia, M. Maadooliat, R.K. Woods, S.H. Audi, B.J. Tefft, C. Chiastra, E.S.H. Ibrahim, J.F. LaDisa, Aortic Remodeling Kinetics in Response to Coarctation-Induced Mechanical Perturbations, Biomedicines 11 (2023). 10.3390/biomedicines11071817.

[66] A. Menon, D.C. Wendell, H. Wang, T.J. Eddinger, J.M. Toth, R.J. Dholakia, P.M. Larsen, E.S. Jensen, J.F. LaDisa, A coupled experimental and computational approach to quantify deleterious hemodynamics, vascular alterations, and mechanisms of long-term morbidity in response to aortic coarctation, J Pharmacol Toxicol Methods 65 (2012) 18–28. 10.1016/j.vascn.2011.10.003.

[67] S. Dutta, P. Sengupta, Men and mice: Relating their ages, Life Sci 152 (2016) 244–248. 10.1016/j.lfs.2015.10.025.

[68] E.C. Davis, Histochemistry Stability of elastin in the developing mouse a quantitative radioautographic study, 1993.

[69] J.E. Wagenseil, R.P. Mecham, Vascular Extracellular Matrix and Arterial Mechanics, Physiol Rev 89 (2009) 957–989. 10.1152/physrev.00041.2008.

[70] L.C.Y. Wong, B.L. Langille, Developmental Remodeling of the Internal Elastic Lamina of Rabbit Arteries Effect of Blood Flow, 1996. http://ahajournals.org.

[71] G. Cardoso, M. Abecasis, R. Anjos, M. Marques, G. Koukoulis, C. Aguiar, J.P. Neves, Aortic coarctation repair in the adult, J Card Surg 29 (2014) 512–518. 10.1111/jocs.12367.

[72] A.B. Bhatt, M.R. Lantin-Hermoso, C.J. Daniels, R. Jaquiss, B.J. Landis, B.S. Marino, R.H. Rathod, R.N. Vincent, B.B. Keller, J. Villafane, Isolated Coarctation of the Aorta: Current Concepts and Perspectives, Front Cardiovasc Med 9 (2022). 10.3389/fcvm.2022.817866.

[73] C.M. Vrbin, Parametric or nonparametric statistical tests: Considerations when choosing the most appropriate option for your data, Cytopathology 33 (2022) 663–667. 10.1111/cyt.13174.

[74] K.J. Bubb, A.A. Aubdool, A.J. Moyes, S. Lewis, J.P. Drayton, O. Tang, V. Mehta, I.C. Zachary, D.J. Abraham, J. Tsui, A.J. Hobbs, Endothelial C-Type Natriuretic Peptide Is a Critical Regulator of Angiogenesis and Vascular Remodeling, Circulation 139 (2019) 1612–1628. 10.1161/CIRCULATIONAHA.118.036344.

