## Supplement for "Structural and Mechanical Analysis of Treated and Untreated Aortic Coarctation in a Growing Porcine Model"

### SUPPLEMENTARY MATERIAL

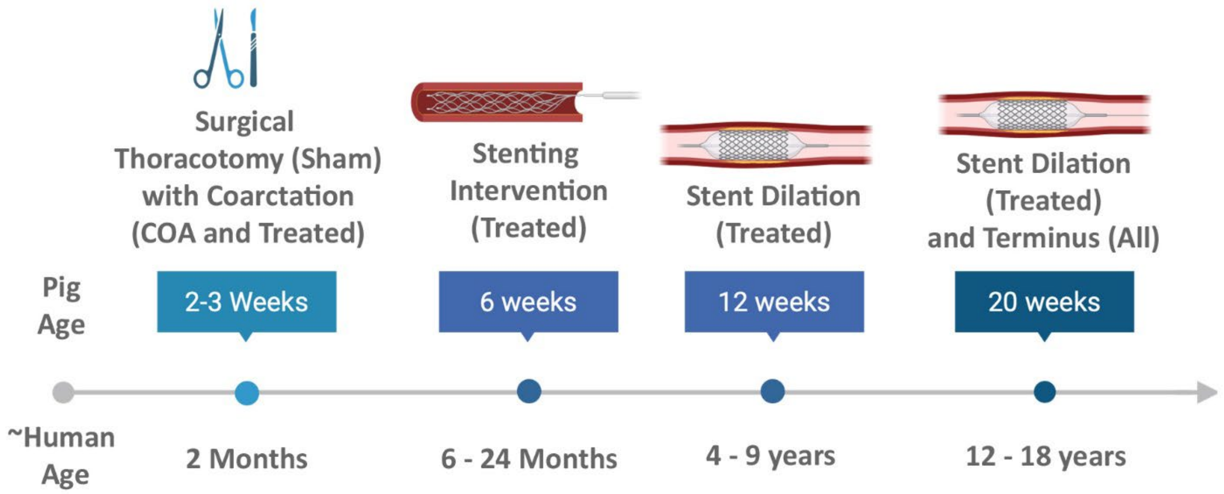


**Supplemental Figure 1:** The experimental timeline began with surgical thoracotomy when pigs were 2 weeks of age and extended to terminal harvest at 20 weeks of age. The timeline also indicates the approximate human age for each timepoint. Created in BioRender. Gober, L. (2025) https://BioRender.com/c2prk0a

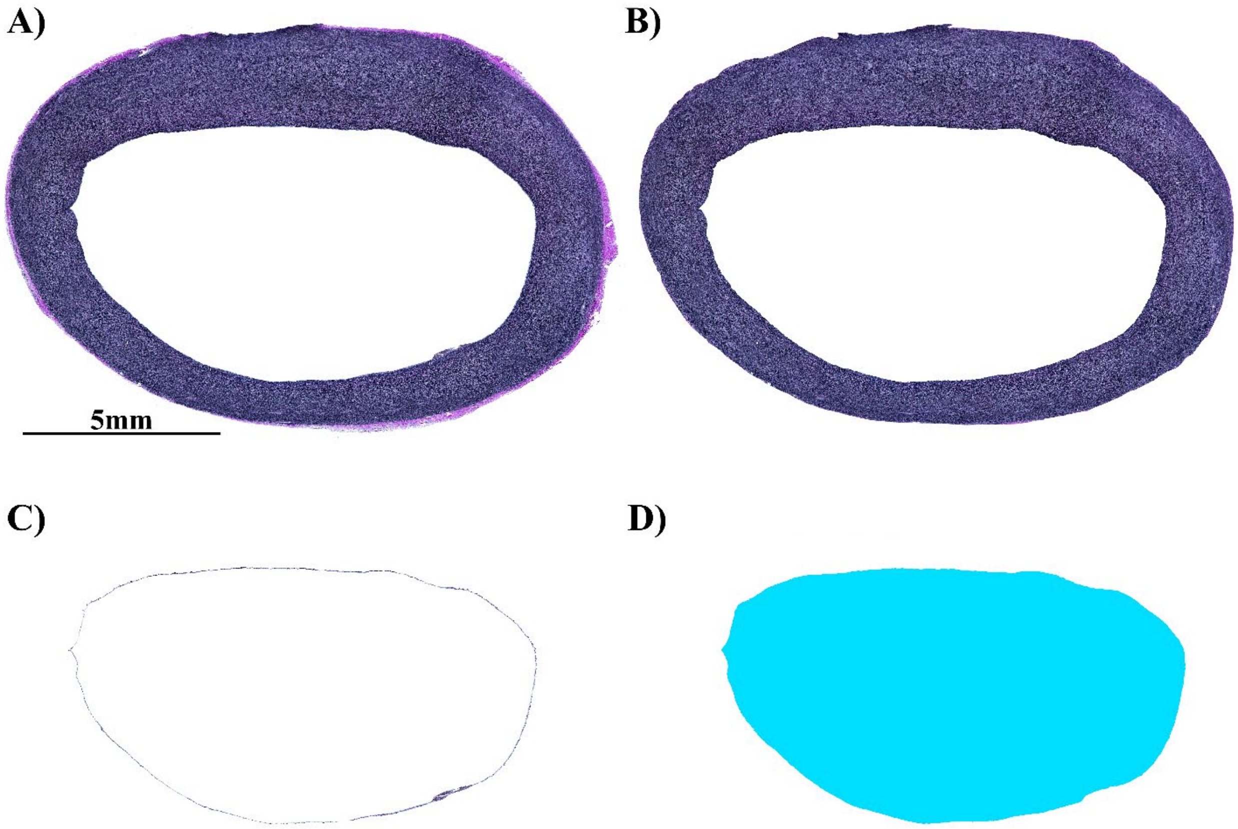


**Supplemental Figure 2:** Representative images showing segmentation of a proximal sham aortic cross-section with A) initial raw image, B) media only, C) intima only, and D) lumen.

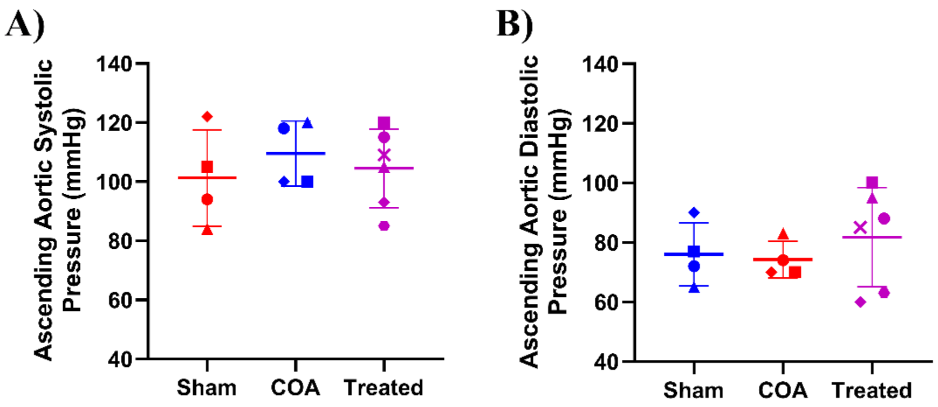


**Supplemental Figure 3**: (A) Ascending aortic systolic pressure at 20 weeks for the sham, COA control, and treated animals post-dilation. (B) Ascending aortic diastolic pressure at 20 weeks for the sham, COA control, and treated animals post-dilation. There were no statistically significant differences between groups for a Kruskal-Wallis test with Dunn's multiple comparisons testing.

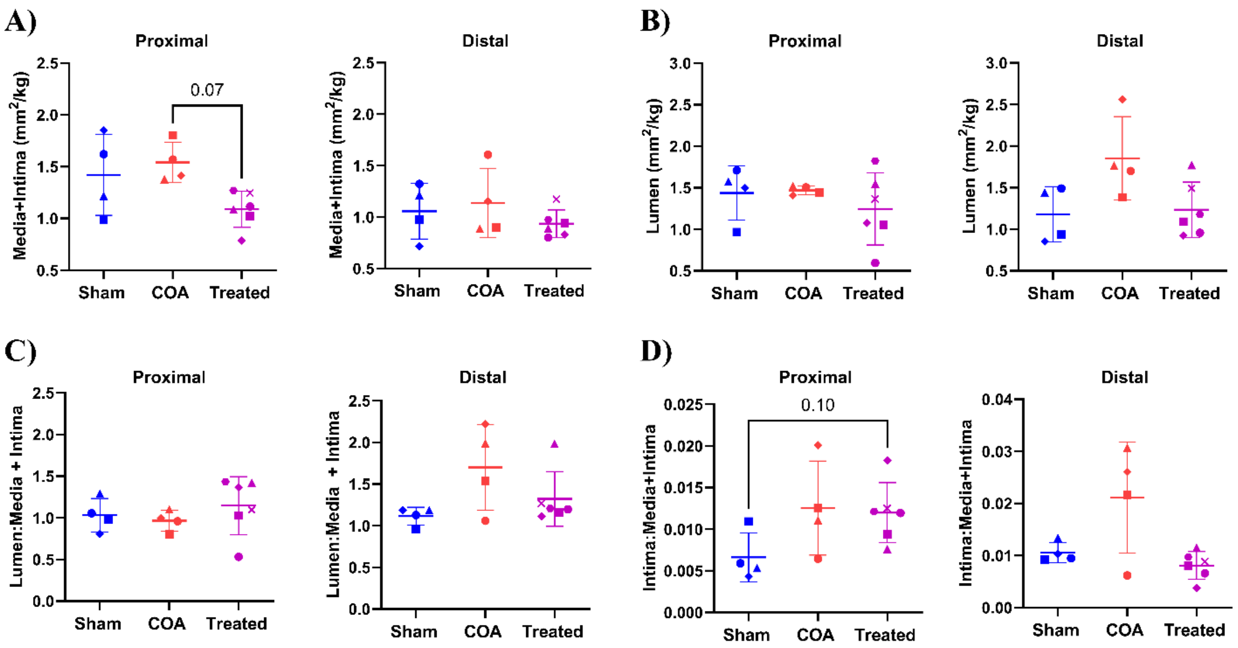


**Supplemental Figure 4**: (A) The tissue area (media + intima) normalized by animal weight immediately proximal and distal to the COA site from COA control and treated animals and comparable regions for the sham animals. (B) The proximal and distal lumen area normalized by weight for the three groups. (C) The proximal and distal ratio of lumen-to-tissue area and (D) the proximal and distal ratio of intima-to-tissue area. Statistical significance was indicated for a Kruskal-Wallis test with Dunn's multiple comparisons testing.

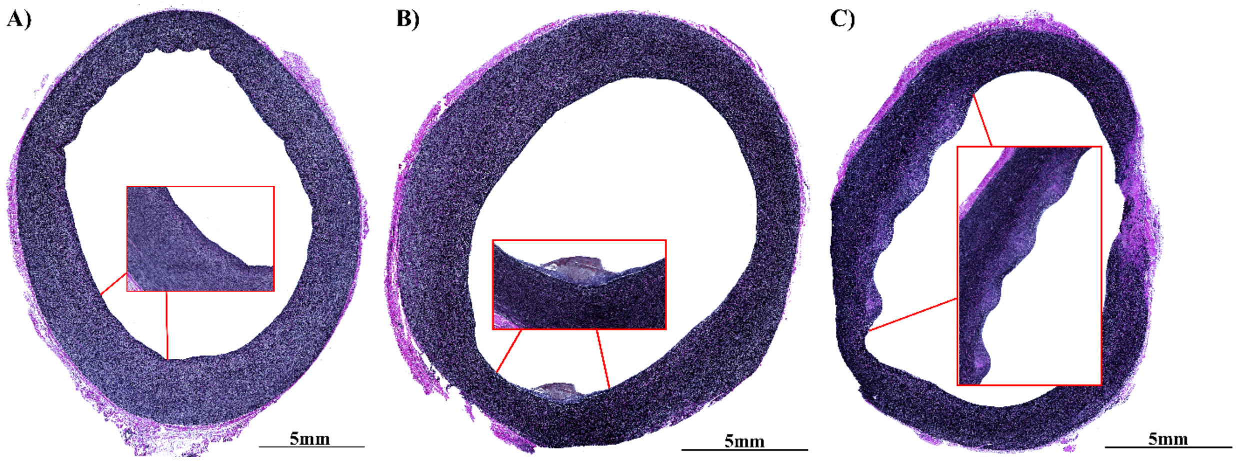


**Supplemental Figure 5:** Representative proximal VVG stained aortic section from a A) sham, B) COA, and C) treated pig. There was marked intimal hyperplasia in sections from both the COA and treated animals.

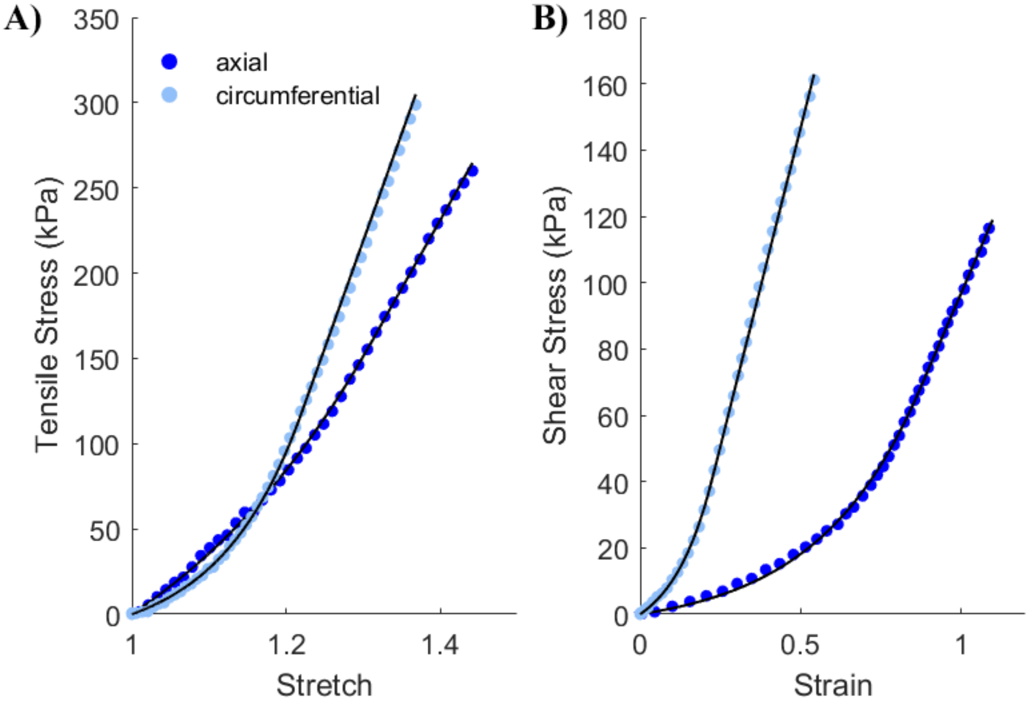


**Supplemental Figure 6:** (A) First Piola-Kirchhoff tensile stress versus grip stretch during uniaxial extension to failure for a representative sham animal with samples oriented along the axial and circumferential directions (dots) as well as the fitted constitutive model (lines). B) First Piola-Kirchhoff shear stress versus Green-Lagrange shear strain during shear lap extension to failure for a representative sham animal with samples oriented along the axial and circumferential directions (dots) as well as the fitted constitutive model (lines).

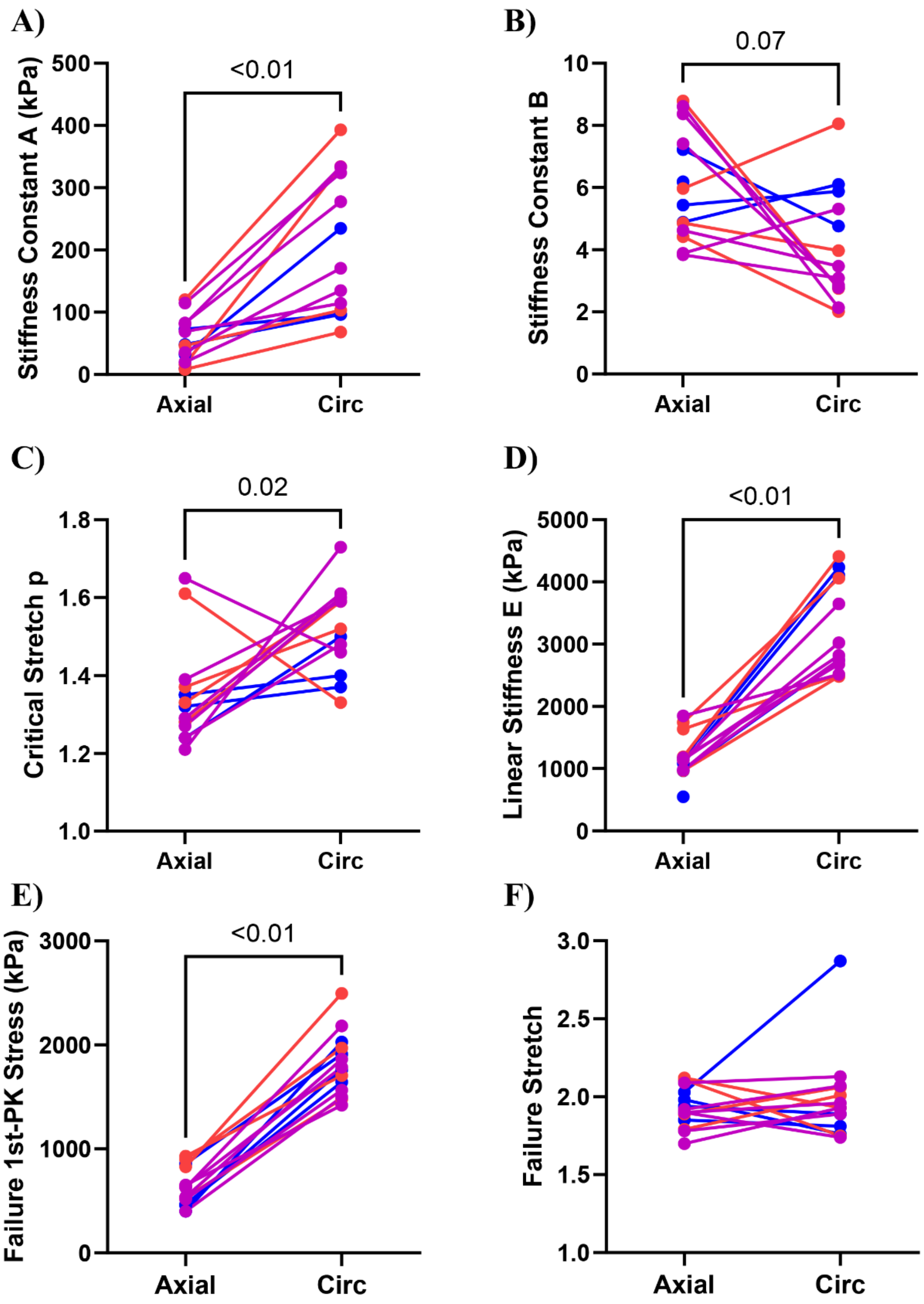


**Supplemental Figure 7**: Pooled toe region stiffness constants $A$ (A) and $B$ (B) for uniaxial samples aligned axially (Axial) and circumferentially (Circ). C) The pooled transition stretch, $p$, between the toe and linear regions and D) the pooled linear stiffness constant, $E$. E) Failure first Piola-Kirchhoff tensile ($1^{st}$-PK) stress and F) failure grip stretch. Statistical significance between the Axial and Circ directions was indicated for a Wilcoxon matched-pairs signed rank test. The data was pooled since there were no statistically significant differences between groups in any metric or its anisotropy according to Kruskal-Wallis tests.

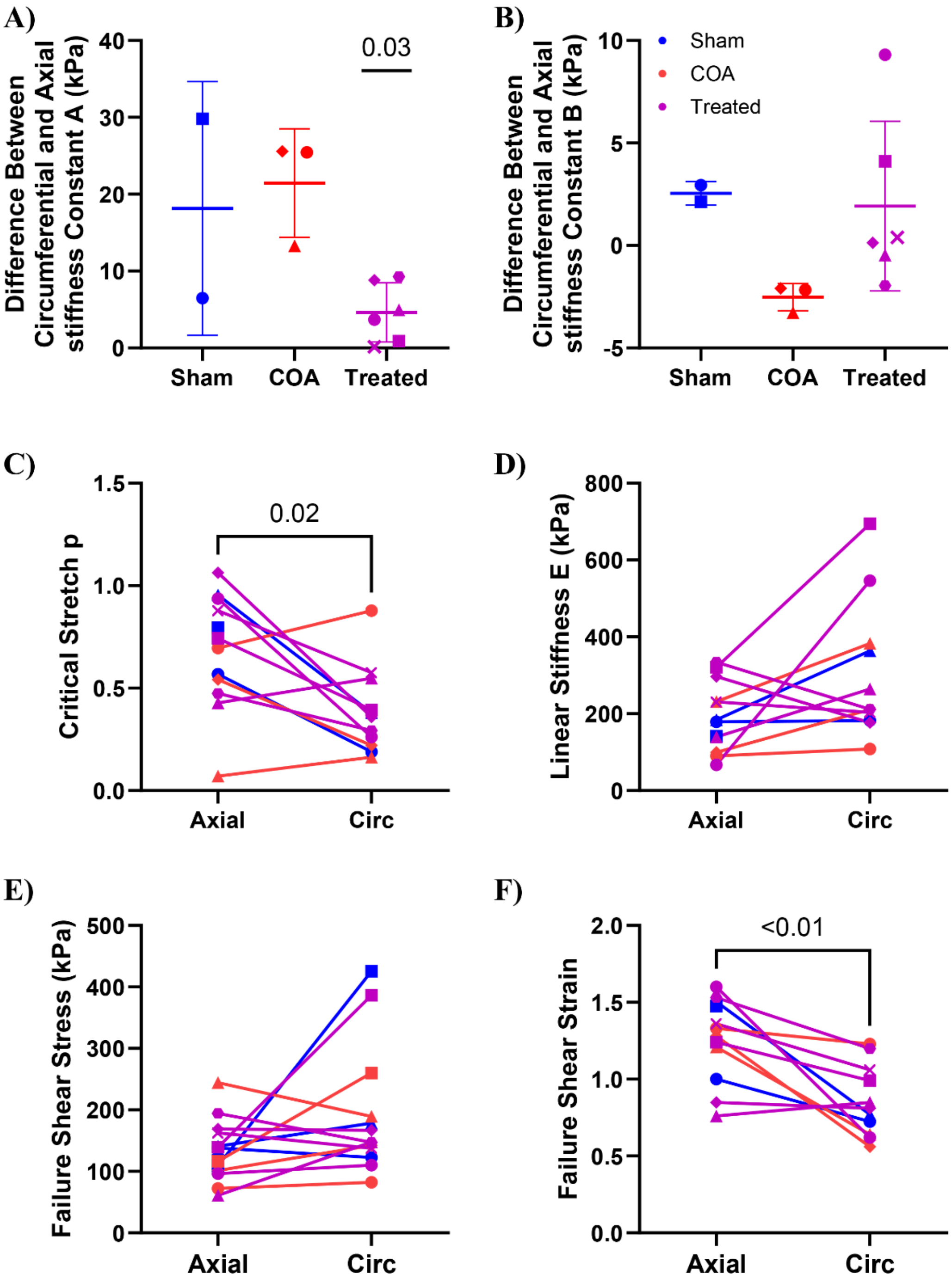


**Supplemental Figure 8**: Anisotropy in the toe region stiffness constants $A$ (A) and $B$ (B) for shear testing were significantly different between groups according to a Kruskal-Wallis test. Therefore, group-wise statistical significance between the circumferential (Circ) and axial (Axial) directions were determined via Wilcoxon matched-pairs signed rank tests. For all other metrics data was pooled since there were no statistically significant differences between groups in anisotropy according to Kruskal-Wallis tests. Pooled transition stretch, $p$, between the toe and linear regions (C),) the pooled linear stiffness constant, $E$, E) pooled failure first Piola-Kirchhoff shear stress, and F) pooled failure Green-Lagrange shear strain. Statistical significance between the Axial and Circ directions was indicated for a Wilcoxon matched-pairs signed rank test.

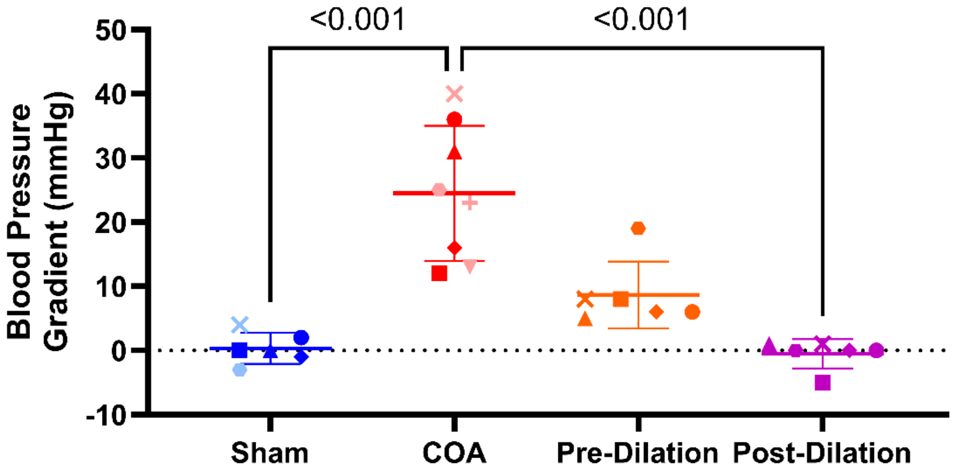


**Supplemental Figure 9**: The peak-to-peak systolic blood pressure gradient across the COA site for the larger COA control group (n = 8) was significantly higher than the pressure gradient across the same anatomic location for the larger sham group (n = 6) at 20 weeks. It was also significantly higher that of the treated group (n = 6) following stent dilation. Statistical significance was indicated for a Kruskal-Wallis test with Dunn's multiple comparisons testing. Animals not included in the original analysis were indicated by light blue and light red symbols.

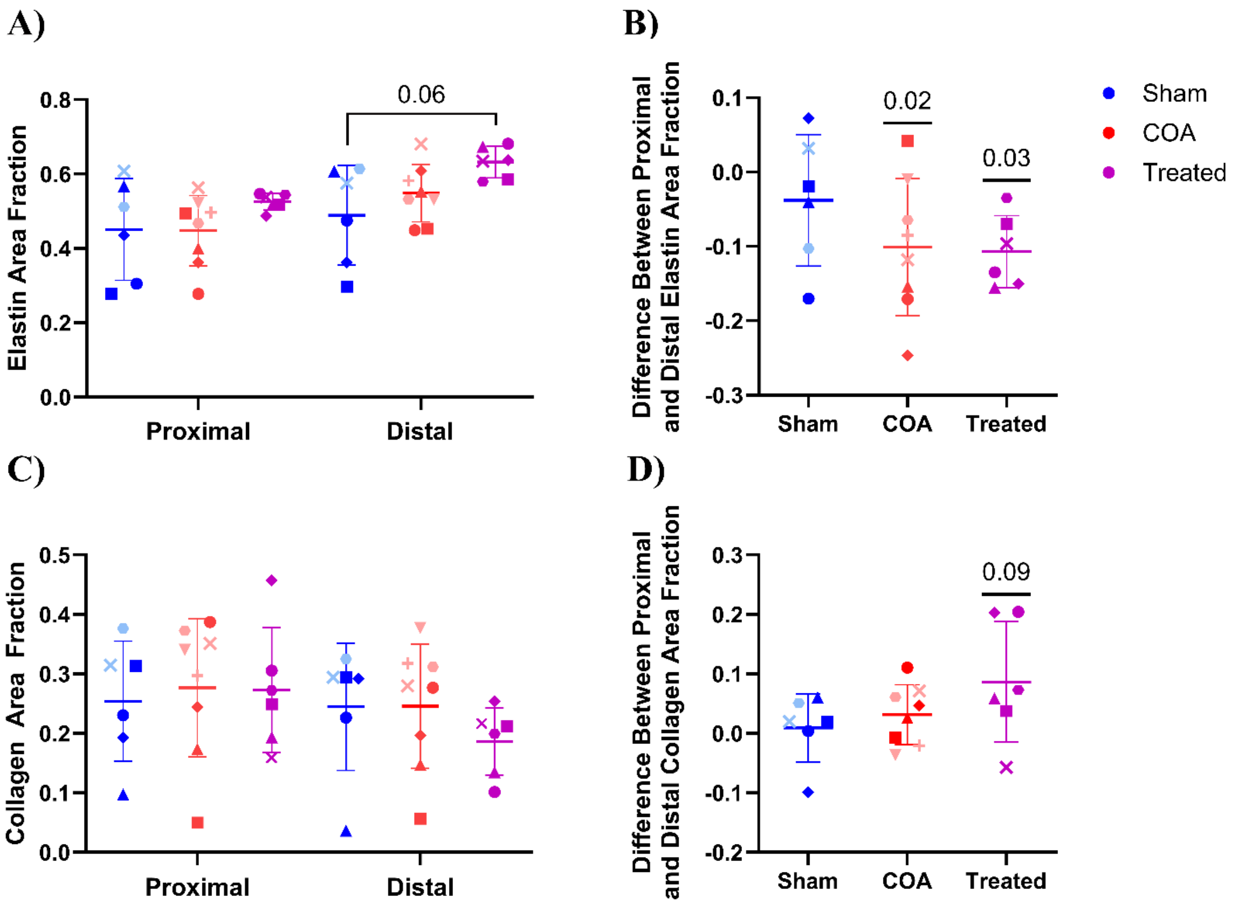


**Supplemental Figure 10:** A) Elastin and C) collagen area fractions computed for aortic cross-sections located in the descending aorta immediately proximal and distal to the COA site from the larger COA control group (n = 8) and the treated group (n = 6) as well as comparable locations for the larger sham group (n = 6). Treated group samples taken distally to the COA site exhibited nearly significantly more elastin than distal sham samples according to Dunn's multiple comparison testing following a Kruskal-Wallis tests. There were no significant differences between groups in the location change (i.e. proximal minus distal) in B) elastin or D) collagen area fraction. However, a one sample Wilcoxon test indicated that in the treated animals there was a significantly lower elastin area fraction proximally and a nearly significantly higher collagen area fraction proximally. A one sample Wilcoxon test also indicated that the larger COA control group exhibited a significantly lower elastin area fraction proximally. Animals not included in the original analysis were indicated by light blue and light red symbols.

**Supplemental Table 1: The number of samples and animals included for uniaxial, shear, and peel testing analyses**

| | Direction | Number of Animals | | | Number of Samples | | |
|---|---|---|---|---|---|---|---|
| | | Sham | COA | Treated | Sham | COA | Treated |
| **Uniaxial Tests** | | | | | | | |
| Pre-Failure Parameters (A, B, p, and E) | Axial | 4 | 4 | 6 | 10 | 12 | 17 |
| | Circ | 3 | 4 | 6 | 9 | 12 | 22 |
| Failure Stretch | Axial | 4 | 4 | 6 | 10 | 12 | 17 |
| | Circ | 4 | 4 | 6 | 10 | 12 | 22 |
| Failure Tensile Stress | Axial | 4 | 4 | 6 | 10 | 12 | 17 |
| | Circ | 4 | 4 | 6 | 14 | 12 | 22 |
| **Shear Lap Tests** | | | | | | | |
| Pre-Failure Parameters (A, B, p, and E) | Axial | 3 | 3 | 6 | 5 | 5 | 13 |
| | Circ | 2 | 3 | 6 | 4 | 5 | 13 |
| Failure Stretch | Axial | 3 | 3 | 6 | 6 | 5 | 13 |
| | Circ | 2 | 3 | 6 | 4 | 5 | 13 |
| Failure Tensile Stress | Axial | 4 | 4 | 6 | 8 | 8 | 14 |
| | Circ | 3 | 4 | 6 | 6 | 8 | 13 |
| **Peel Tests** | | | | | | | |
| Peel Tension | Axial | 4 | 4 | 6 | 9 | 8 | 13 |
| | Circ | 4 | 4 | 6 | 8 | 9 | 15 |
